# Sugars dominate the seagrass rhizosphere

**DOI:** 10.1101/797522

**Authors:** E. Maggie Sogin, Dolma Michellod, Harald Gruber-Vodicka, Patric Bourceau, Benedikt Geier, Dimitri V. Meier, Michael Seidel, Soeren Ahmerkamp, Sina Schorn, Grace D’Angelo, Gabriele Procaccini, Nicole Dubilier, Manuel Liebeke

## Abstract

Seagrasses are one of the most efficient sinks of carbon dioxide on Earth^1^: They bury carbon 35 times faster than tropical rainforests on a per unit area basis^2^. While we know that carbon sequestration in terrestrial plants is intimately linked to the microorganisms living in their soils^3–6^, the interactions of seagrasses with their rhizospheres are poorly understood. We show that three seagrass species from two oceans excrete simple sugars, mainly sucrose, into their rhizosphere that accumulate to over 200 µM. Such high concentrations are at least 80 times higher than previously observed in the ocean, and surprising, as sugars are quickly consumed by microorganisms. In situ analyses and incubation experiments indicated that phenolic compounds from the seagrass inhibited microbial consumption of sucrose. Metagenomic and metatranscriptomic analyses of the microbial communities in the seagrass rhizosphere revealed that many members had the genes for degrading sucrose, but these were only expressed by a few specialists that also expressed genes for degrading phenolics. Our results explain why sucrose accumulates under seagrass meadows, where it comprises as much as 40% of the dissolved organic carbon. Destruction of extant seagrass canopies would allow sediment microorganisms to consume the tremendous deposits of sucrose buried underneath their meadows, thereby releasing large amounts of CO2 into the oceans and atmosphere.

## Main

Seagrasses are marine angiosperms that represent the foundation species of valuable ecosystems widespread along coastal regions of all continents except Antarctica^7^. As ecosystem engineers, seagrasses provide important services to humans. For example, seagrass meadows are important habitats for fisheries, stabilize the sea floor, remove pollutants, and drive biogeochemical cycling^8^. Like their terrestrial relatives, seagrasses play an import role in sequestering carbon from the atmosphere by incorporating it into their tissues and burying it as organic matter in their sediments^1^. Seagrasses also excrete high concentrations of dissolved organic carbon (DOC) into the environment^9^, a reactive carbon source^10, 11^ that is assumed to be quickly metabolized by the microorganisms in the sediments surrounding seagrass rhizomes and roots.

On land, angiosperms excrete up to 30% of their primary production as organic compounds to their soils to attract beneficial microbial partners, defend themselves from pathogens, and to communicate with other individuals^12^. These root exudates feed complex microbial food webs that drive the long-term storage of organic carbon^3^. Much less is known about how seagrasses interact with the microbial communities in their sediments^13, 14^. Here, we show that seagrass sediments are veritable sweet spots in the sea that contain surprisingly high concentrations of simple sugars, primarily in the form of sucrose. Our results indicate that microbial consumption of these reactive sources of organic carbon is inhibited by the presence of phenolic compounds in the seagrass rhizosphere.

### Sediments underneath seagrass meadows are rich in simple sugars

Our metabolomic analyses of porewater chemistry underneath a meadow of the endemic Mediterranean seagrass, *Posidonia oceanica,* unexpectedly revealed high concentrations of sugars in the high µM range (> 200 µM), with concentrations exceeding the detection limits of our mass spectrometer. To determine if sugars occurred in other seagrass meadows, we extended our metabolomic analyses to three other seagrass species from the Caribbean (*<u>Thalassia testudinum</u>* and *Syringodium filiforme*) and Baltic Sea (*Zostera marina*) that belong to three phylogenetically distinct lineages of marine angiosperms. We consistently measured sugars in the porewater sediments underneath all three seagrass species, with similarly high concentrations above 200 µM in the Caribbean meadows as in the Mediterranean, and concentrations in the low µM range underneath meadows in the Baltic Sea (**Figure 1 a, b; Table S1, Supporting Information**). The primary sugar in the porewater metabolome of the Mediterranean and Caribbean seagrasses was the disaccharide sucrose, which has been shown to be the most abundant sugar in the tissues of many seagrass species, including all four species studied here^15, 16^. Given that seagrasses occupy between 300,000-600,000 km^2^ of coastal regions^1^, we conservatively estimated a global pool of 0.56 - 1.12 Tg of sucrose in the upper 30 cm of seagrass sediments. The sucrose concentrations we measured underneath seagrasses are as much as 80 times higher than previously measured dissolved total carbohydrate pools from marine sediments and 2000 fold higher than free sugar concentrations in the open ocean^17, 18^. Our observations represent a paradox in microbial ecology, as it is widely known that most microorganisms, with the exception of some that live in nutrient limited soil ecosystems, quickly consume sugars in their immediate environment^19–21^. Until now, the only known microbial habitats that contain similarly high concentrations of simple sugars include saps and fruit nectars, rhizospheres of agroecosystems including sugar beets, and biofilms on mosses and marine macroalgae^22, 23^.

**Figure 1.**
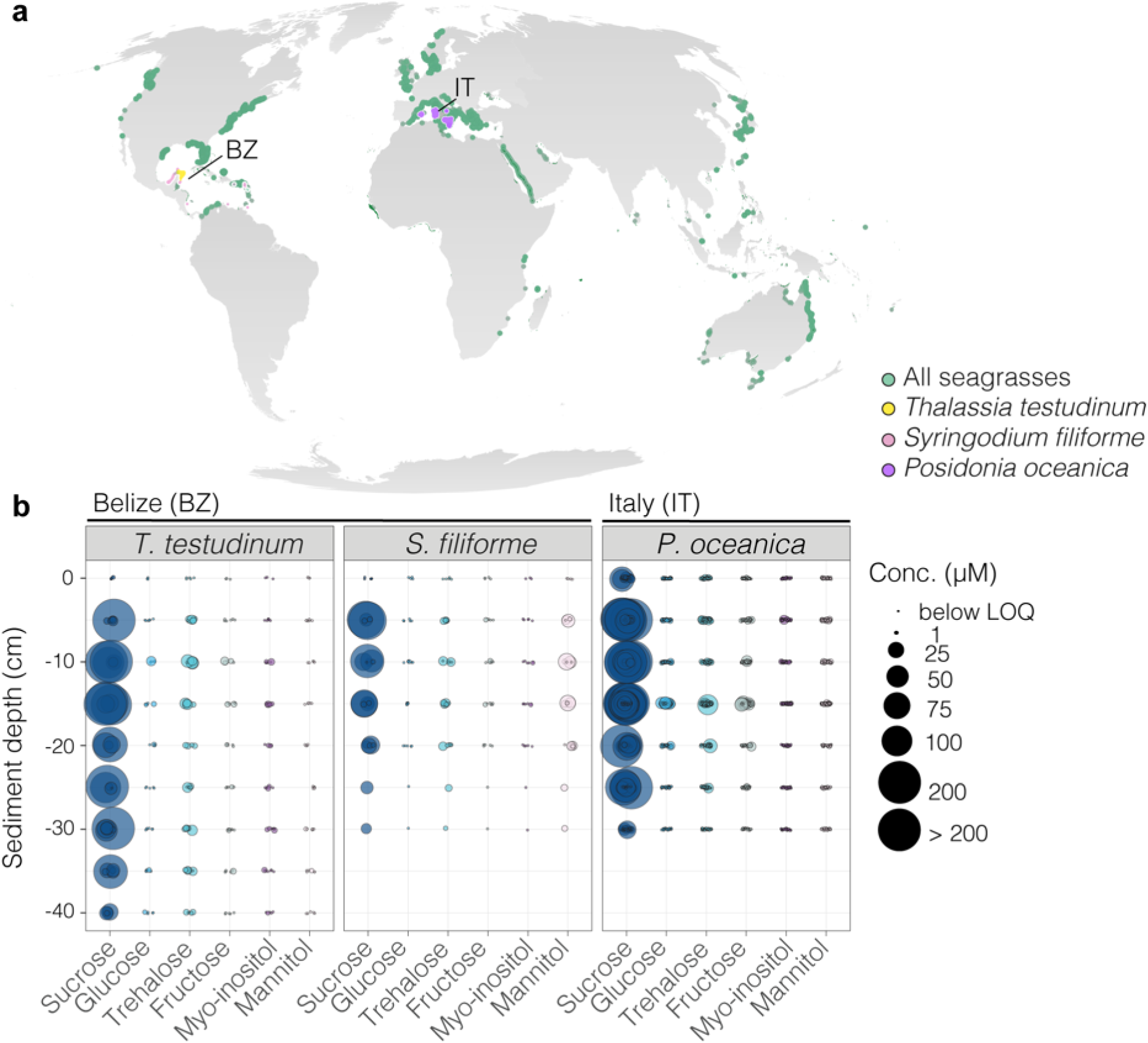
Sediment porewaters underneath seagrass meadows worldwide contain high concentrations of sugars. **a**, Seagrasses occur in coastal waters worldwide (data accessed from UN Environmental World Conservation Monitoring Center on March 21, 2020, http://data.unep-wcmc.org/datasets/7). *Syringodium filiforme* (pink), *Thalassia testudinum* (yellow) in Belize (BZ) and *Posidonia oceanica* (purple) in Italy (IT) were the dominant species at both sampling sites in this study. **b,** Bubble plots show porewaters collected underneath seagrass meadows were rich in sugars in Belize and Italy. The size of each point represents the concentration of each sugar in porewaters from the sampled sediment depths (**Table S1**). For each seagrass meadow, between 6 (*S. filiforme, T. testudinum*) and 14 (*P. oceanica*) individual profiles were sampled (each profile was subsampled in 5 cm steps from 0 – 30 cm sediment depth). To facilitate visualization, samples where sugars were not detected are not plotted, samples in which sugars were detected but below reliable quantification limits are sized to the smallest point and colored based on type of sugar, while values above our quantification limit of 200 µM are sized to the largest point, but were considerably higher. LOQ = limit of quantification.

To understand sugar distribution across spatial scales, we collected 570 metabolomic profiles from sediment porewater underneath, 1 m and 20 m away from a *P. oceanica* meadow in the Mediterranean (i.e., In, Edge, Out; **Figure 2a**). Non-targeted and targeted analyses revealed that sediments underneath *P. oceanica* contained significantly higher concentrations of sucrose, glucose, trehalose, myo-inositol, and mannitol compared to the sites away from the seagrass (**Figure 2b; Extended Data 1; Table S2**). Bulk measurements of DOC were at least 2.6 times higher inside the meadow than at the edge or outside (**Figure S1a; Table S2**).

**Figure 2.**
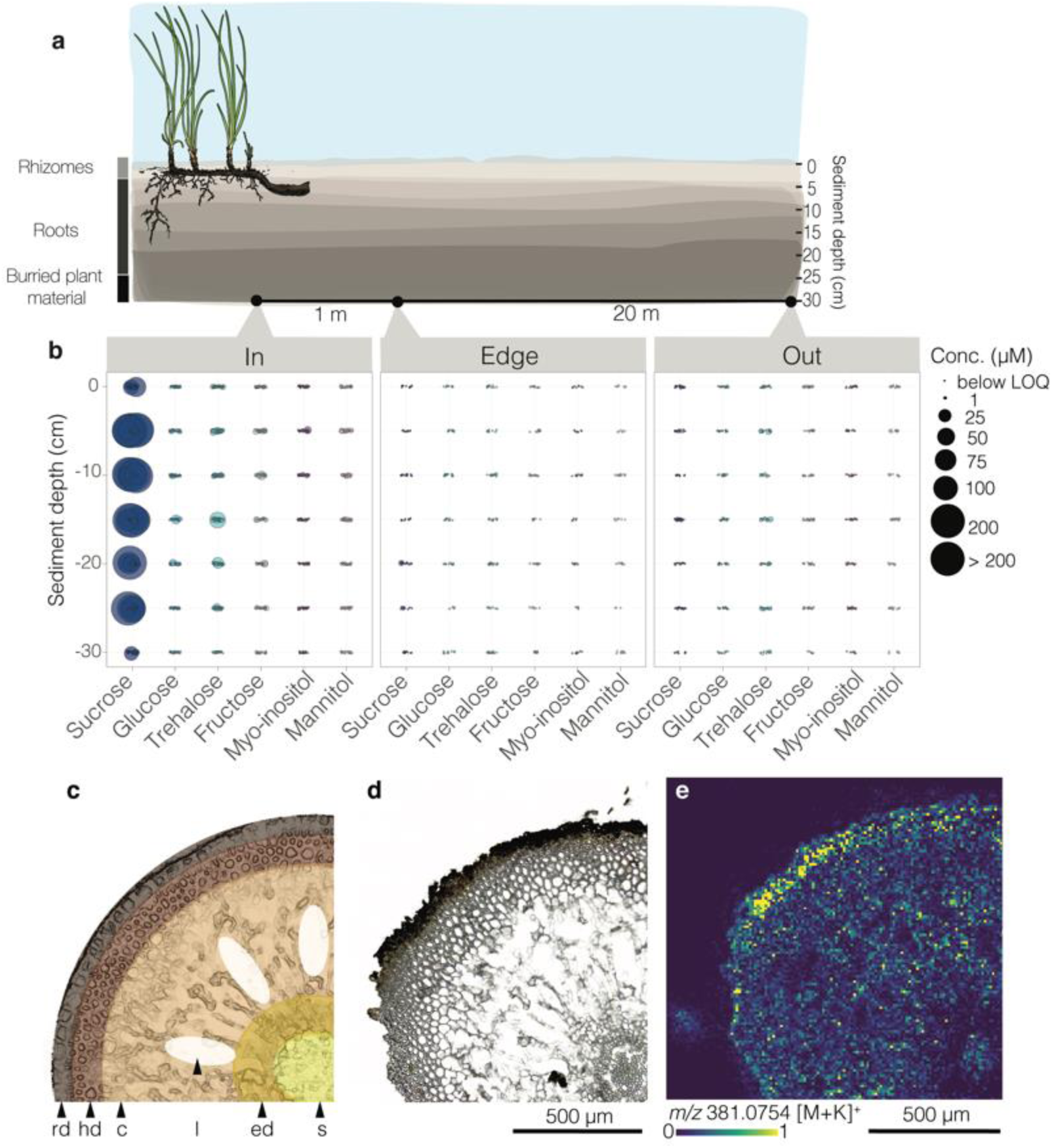
Sugars were more abundant underneath seagrass meadows than in non-vegetated sediments. **a**, Sediment porewater profiles were collected underneath, 1 m and 20 m away from a Mediterranean *P. oceanica* meadow off the Island of Elba, Italy. Porewater profiles consisted of samples collected every 5 cm from 0 to 30 cm sediment depth. **b,** Bubble plots show porewater sugar concentrations across sediment depth. Sugar concentrations were significantly higher inside the meadow than outside or at the edge (two-way ANOVA comparing sugar concentrations as a function of sediment depth and location; model results and significance values reported in **Table S2**). Sugar concentrations above 0 but below reliable quantification are sized to the smallest point, while values above 200 µM (upper detection limit) are sized to the largest point. **c,d,** Schematic, light microscopy and **e,** matrix-assisted laser desorption/ionization mass spectrometry images from a *P. oceanica* root collected from a meadow in the Mediterranean off the Island of Elba, Italy. Displayed is a quarter of a root cross section. In **e,** the molecular ion distribution of disaccharides (C_12_H_22_O_11_, potassium adduct [M+K]^+^, *m/z* 381.0754), such as sucrose, shows the relative abundance was highest in the root rhizodermis. Relative abundances are visualized as a heatmap from low (blue) to high (yellow) intensities of each pixel (10 µm). rd = rhizodermis, hd = hypodermis, c = cortex, l = lacunae, ed = endodermis, s = stele. LOQ = limit of quantification.

Measured sugars contributed up to 40 % of the DOC pool underneath seagrass meadows that form an energy-rich source of carbon that has not yet been described or characterized (**Figure S1b**)^9^. Previous studies showed that seagrasses excrete organic carbon produced from CO_2_ fixation into their sediments^24, 25^, and it has been hypothesized that sucrose is one of the major components of the excreted carbon given its high concentrations in seagrass tissues^15, 16^. However, until now the molecular composition of the organic carbon excreted by seagrasses was not known. Here, we show that sucrose is the major component of seagrass exudates, and in the following, provide evidence linking seagrass primary production to the accumulation of sucrose in its rhizospheres.

Sugar concentrations in Mediterranean *P. oceanica* tissues and sediments were correlated to the availability of light on both seasonal and daily time scales. Our seasonal analyses revealed significantly higher sucrose concentrations in sediment porewaters underneath a *P. oceanica* meadow in April and October than in July (**Figure S2; Table S2**). These observations are consistent with *P. oceanica’s* growing season over the summer months. The observed decline in sucrose concentrations, along with previous observations of decreases over the summer in total organic carbon and bulk carbohydrates in sediments underneath *P. oceanica*^26–28^, suggests that the seagrass diverted these organic carbon resources to support growth during their main productivity season. To understand daily fluctuations of sucrose, we measured sucrose concentrations at discrete sampling time points over 24 hours in both *P. oceanica* tissues and their sediments. Our data indicates that during the day, *P. oceanica* produces sucrose in its leaves as a byproduct of photosynthesis, which is transferred to the roots and rhizomes where it is released into the porewater. Sucrose abundances in *P. oceanica* leaves, but not their rhizomes or roots, were significantly higher at dusk than at all other sampling time points (**Extended Data 2a-d; Table S2**). Changes in sucrose abundances within *P. oceanic*a leaves thus followed the diel ramping up and down of photosynthetic and respiration rates^29–31^. Relatively stable sucrose concentrations in *P. oceanica*’s roots and rhizomes over 24 hours suggest that the regulation of sucrose in these organs is independent of diel rhythms. Within the porewaters underneath the seagrass, sucrose concentrations were significantly higher during the day than at night (**Extended Data 2e; Table S2**). In summary, our data indicates excess sucrose production during peak productivity hours is transferred to the underlying sediments, where it is only partly consumed during the night.

To reveal how sucrose is excreted and distributed in *P. oceanica* roots, we used mass spectrometry imaging (MSI) to visualize the molecular distribution of sugars in the seagrass roots. Mass spectrometry images of cross sections of seagrass roots revealed that sucrose was distributed throughout the root tissues and highly abundant in the rhizodermis, the outermost cell layer of the root (**Figure 2c-e; Figure S3**). This suggests that sucrose is excreted, or passively released to the sediment through the rhizodermis. Both active transport and diffusion are known from terrestrial plants which release exudates, including sugars, through their roots into soils^12, 32^. Our sediment porewater profiles provided further support for our assumption that sucrose enters the rhizosphere through the roots, as we measured the highest sucrose concentrations at the sediment depths where the seagrass roots occurred (**Extended Data 3a, b**; **Figure S4**). Collectively, our results from three seagrass species in the Caribbean and Mediterranean seas showed that seagrasses transfer massive amounts of sucrose to marine sediments through their roots on a daily basis. (For a discussion of explanations for the relatively low concentrations of sugars in porewaters from seagrass in the Baltic Sea, see **Supporting Information.**)

### Sediment chemistry limits microbial respiration of sugars underneath *P. oceanica* meadows

Sucrose and other sugars are among the most favorable and important carbon and energy sources for soil and sediment microorganisms. To understand why seagrass produced sugars are not quickly consumed by sediment microorganisms, we measured in situ oxygen concentrations underneath a Mediterranean *P. oceanica* meadow. We observed a sharp decrease in oxygen concentrations, from 202 µM to < 0.5 µM, within the first 7.5 cm below the sediment surface (**Extended Data 3c**). The observed decrease in oxygen with depth in sediment porewaters suggests that microorganisms below 7.5 cm are not able to use aerobic pathways for sucrose respiration. In the absence of oxygen, microorganisms ferment sugars to organic acids, which a syntrophic community of bacteria and archaea respire to CO_2_ using mainly nitrate and sulfate as electron acceptors. While nitrate is limited in sediments underneath seagrass meadows^33^, sulfate concentrations in Mediterranean porewaters exceed 28 mM and are stable across sites and sediment depths^34^. It is therefore likely that there was sufficient sulfate underneath *P. oceanica* meadows to allow sulfate-reducing microorganisms to metabolize the waste products of sugar fermentation. Coupled with previous observations of little to no sulfide underneath these meadows despite the availability of sulfate^34^, we hypothesize that other biogeochemical factors beyond electron acceptors inhibit the microbial degradation of simple sugars underneath seagrass meadows.

Phenolics represent one class of compounds that could limit the microbial degradation of sucrose under anoxic conditions, as shown for ruminal microbiomes where the presence of phenolics inhibits microbial growth^35^. To test our hypothesis, we analyzed the molecular composition of sediment porewater underneath a Mediterranean *P. oceanica* meadow using ultrahigh-resolution mass spectrometry. We found that between 10 and 16% of all detected molecular formulae consisted of aromatic compounds, including polyphenolics (**Extended Data 4**), which is comparable to ecosystems with strong inputs of terrestrial DOM, such as rivers^36, 37^. Seagrasses, including *P. oceanica,* contain large concentrations of phenolic compounds in their tissues^38, 39^ including their roots (**Extended Data 5**), making the plants and their debris the likely source of phenolics as inhibitory compounds in the seagrass sediments.

To test if the presence of phenolics restricts microbial degradation of sucrose, we isolated phenolic compounds from seagrass tissue and co-introduced this extract with ^13^C-sucrose to meadow sediments under oxic and anoxic conditions. We observed that the presence of seagrass-derived phenolics significantly decreased sucrose respiration rates, and consequently the production of CO_2_ over 10-fold, under anoxic conditions. In contrast under oxic conditions, we saw no significant effect of phenolics on sucrose respiration (**Figure 3a,b; Supporting Information**). This corresponds with our expectation that phenolics limit microbial respiration predominantly under anoxia as previous studies showed that phenolics are quickly degraded under oxic conditions^40^. The degradation of phenolics in the oxic layer under the meadow would allow the sediment microbial community to fully respire sucrose, while under anoxic conditions, high concentrations of phenolics likely inhibit microbial consumption of sucrose, and sugars accumulate under the seagrass meadows.

**Figure 3.**
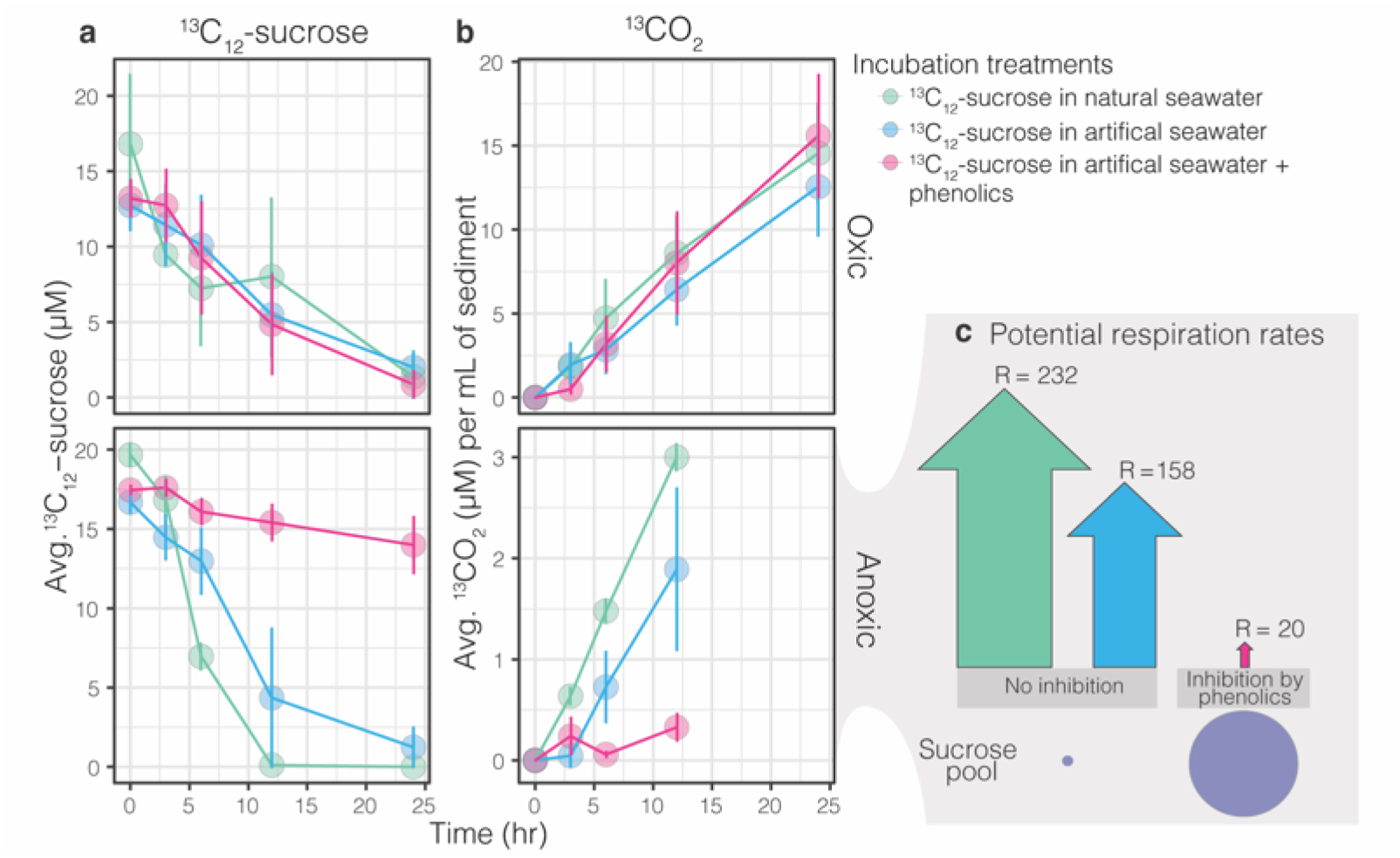
Phenolic compounds inhibit the degradation of sucrose. Sediments collected from inside a Mediterranean *P. oceanica* meadow off the Island of Elba, Italy consumed **a,** ^13^C_12_-sucrose and produced **b,** ^13^CO_2_ over the course of oxic and anoxic incubation experiments conducted over 24 hours. Points represent treatment means, error bars are the standard error of the mean. Under oxic conditions, the production rate of ^13^CO_2_ from ^13^C_12_ -sucrose was similar across treatments and at least 2.5 times higher than rates observed under anoxia (note different y-axis scales). However, in the anoxic incubations, the addition of phenolics extracted from *P. oceanica* tissues inhibited the microbial degradation of sucrose to CO_2_. **c**, The estimated potential respiration rates under anoxia show that the production of ^13^CO_2_ from sucrose in the presence of phenolic compounds is at least 8 times lower than that of natural or artificial seawater condition (**Table S3**). The size of the arrows is proportional to the potential rate of CO_2_ released from sucrose in mmol(C) m^-2^d^-1^.

Our incubation experiments show that the microbial community underneath seagrass meadows has the metabolic ability to degrade sucrose, but this potential is significantly reduced by the presence of phenolics in their environment. Our work, together with research on plant exudation in terrestrial rhizospheres, indicates both marine and land plants excrete sugars and phenolic compounds to their soils^41, 42^. However, in terrestrial rhizospheres where there is sufficient oxygen, these phenolic compounds are degraded^43^. This removes the inhibitory effect of most phenolic compounds on microbial metabolism, so that microbial communities are able to consume plant produced sugars^44^. In contrast, in the marine rhizosphere, only the first few cm underneath the meadow are oxic, while most of the sediment underneath the meadows is anoxic. In these anoxic sediments, seagrass produced phenolics, which enter the sediments either through plant secretion or leaching from decaying peat, are not quickly oxidized and thus accumulate. These phenolics limit the microbial degradation of sucrose, allowing sugars to also accumulate in the seagrass rhizosphere. Thus, the presence of phenolics and the lack of oxygen underneath seagrass meadows delays the input of reactive, organic carbon into microbial respiration and ultimately the microbial loop, allowing sucrose to accumulate.

### Most microbial members of the seagrass rhizosphere do not degrade sucrose

Plant carbon deposition to terrestrial soils serves as a chemo-attractant for beneficial microorganisms, including arbuscular mycorrhizas, and beneficial bacteria and archaea^12^. We hypothesized that seagrass deposition of sucrose, phenolics, and other compounds into their sediments structures the composition and function of the marine rhizosphere.

Using metagenomics and full-length 16S rRNA amplicon sequencing, we show that the sediments underneath seagrass meadows contained a distinct community of microorganisms (**Figure 4**). We extracted genomic DNA and RNA from the 10-15 cm layer of sediment push cores collected inside, 1 m and 20 m outside a Mediterranean *P. oceanica* meadow. We explored the microbial communities at 10-15 cm because this sediment depth contained the highest sucrose concentrations in porewater profiles and was the average depth of *P. oceanica* root penetration. Using principal coordinate analyses, samples clustered according to their collection site, based on the taxonomic composition of both metagenomic and full length 16S rRNA sequencing libraries (**Figure 4a-b**). Our PacBio amplicon sequencing of the full length 16S rRNA gene further revealed that bacterial communities were highly diverse at all three collection sites, mirroring known complexities in terrestrial habitats that contain thousands of taxa per gram of soil^4^. However, we observed fewer amplicon sequence variants inside than outside of the meadow (**Figure 4c**). These data indicate the presence of the seagrass rhizosphere not only selects for specific taxa, but also limits the diversity of the community. On land, plants secrete compounds to their rhizospheres, creating a habitat that shapes the community composition of their soil taxa^45^, and results in a specialized microbial community with lower diversity than neighboring unvegetated soils^6, 46^.

**Figure 4.**
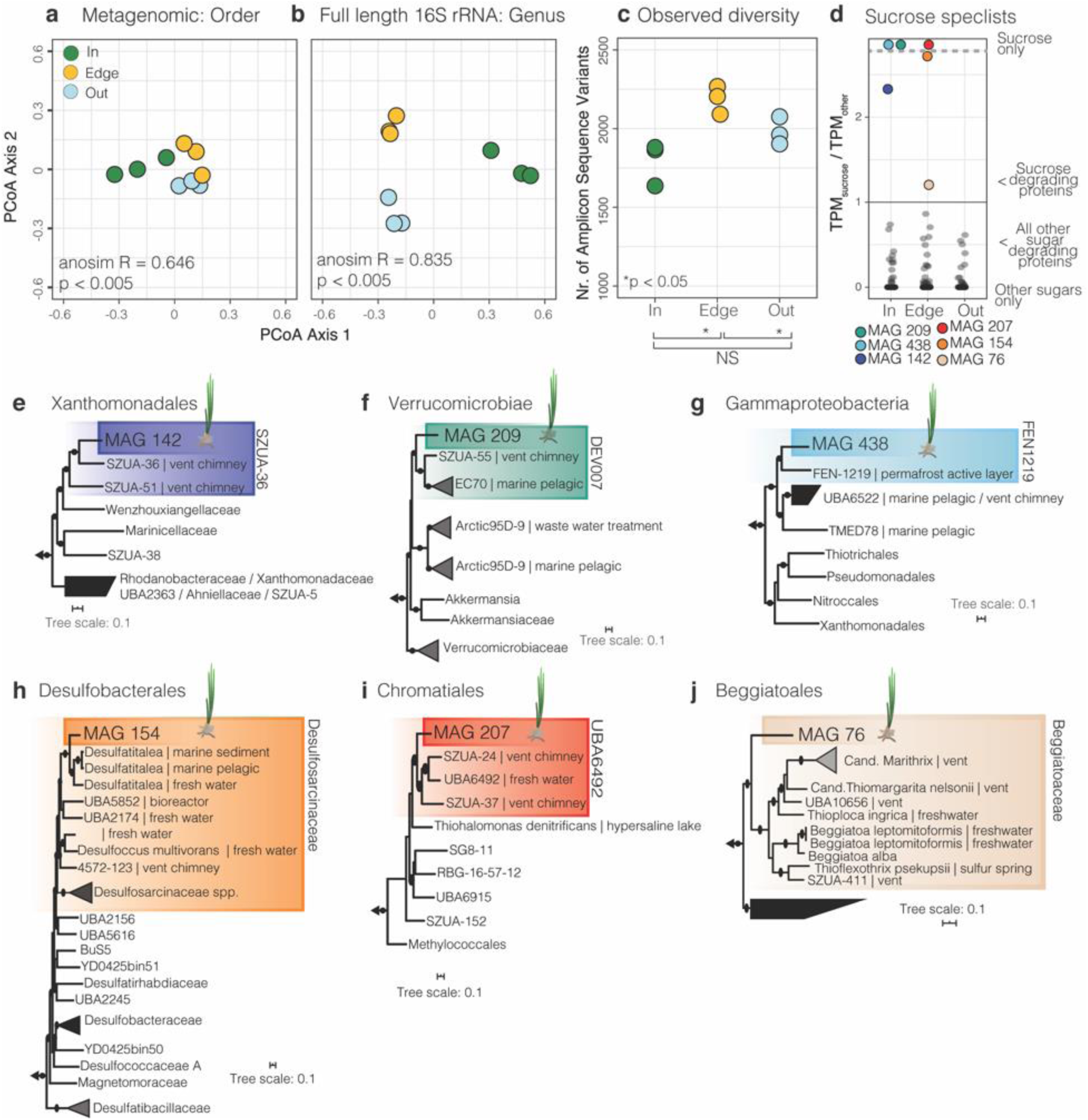
Microbial community composition and metabolism were specific to seagrass sediments. PCoA analyses show that the taxonomic composition of microbial communities underneath (In) and away (Edge and Out) from a Mediterranean *P. oceanica* meadow off Elba, Italy were significantly different based on **a,** bulk metagenomics (order level, ANOSIM R^2^= 0.65, *p-value* < 0.007) and **b,** full-length 16S rRNA amplicon sequencing (genus level, ANOSIM R^2^= 0.78, *p-value* < 0.004). **c,** The total number of Amplicon Sequence Variants (ASV) was significantly lower in samples collected inside and outside the meadow than at the edge (*M*_inside_ = 1793 ± 79 SEM; *M*_edge_ = 2224 ± 23 SEM; *M*_out_ = 1983 ± 51 SEM) **d,** Boxplots show the accumulative expression ratio in each MAG between glycoside hydrolases (GH) predicted to degrade sucrose compared to other sugars (overlying points). Sucrose specialists (colored points) are defined as having higher transcript per million base pair values (TPMs) for sucrose degradation compared to all other GH enzymes (TMP_sucrose_ \ TPM_other_ > 1). TPMs represent the total expression across each site-specific library. The majority of MAGs across sites were sugar generalists, while sucrose specialists were only found in three MAGs inside and three at the edge of the meadow. Phylogenomic trees for 6 sucrose specialists: **e,** MAG 438, **f,** MAG 142, **g,** MAG 209, **h,** Mag 154, **i,** MAG 207 **j,** MAG 76. Colored boxes show members that grouped within the same family clade. For the relative abundance of each of the six sucrose specialists across habitats see **Figure S8**.

Microbial genomes recovered from our metagenomic analyses revealed that taxa in the *P. oceanica* rhizosphere were limited in their ability to degrade sucrose in spite of the high abundance of this simple sugar in their environment (for bioinformatic methods see **Figure S5**). We recovered 182 metagenomic assembled genomes (MAGs), 125 of which were of medium draft quality (**Table S4; Table S5; Figure S6**)^47^, and focused our metagenomic and metatranscriptomic analyses on genes used in microbial sucrose metabolism. While 80% of the recovered MAGs contained genes that degrade sucrose, including sucrose phosphorylases and sucrose-6 phosphate hydrolases, these genes were expressed in only 59% of the MAGs. To analyze if sucrose degradation was central to the metabolism of these MAGs, we compared the proportion of transcripts for sucrose degradation to all expressed orthologs in transcripts per million base pairs (TPM) within a MAG. The expression of sucrose degradation genes in most MAGs, with a few exceptions, were not among the top 10% of all expressed orthologs at all sampling sites, including those directly underneath the meadow with the highest sucrose concentrations (**Figure S7**).

Despite the relatively low expression of sucrose degradation genes across most MAGs, we identified six MAGs across sampling sites where expression profiles showed a higher proportion of sucrose degradation genes compared to genes encoding the degradation of other sugars (TPM_sucrose_ / TPM_other_ > 1; **Figure 4d**; **Extended Figure 6**; **Supporting Information**). This indicates that a select set of sediment taxa preferentially metabolized sucrose over other sugars. The three sucrose specialists, which used sucrose inside the meadow, were undescribed members of the Xanthomonadales (MAG 142), Verrucomicrobiales (MAG 209), and environmental gammaproteobacterial lineage UBA6522 (MAG 438)(**Figure 4e-g**). The sucrose specialists, which used sucrose at the edge of the meadow, belonged to the Desulfobacterales (MAG 154), Thiohalomonadales (MAG 207), and Beggiatoales (MAG 76)(**Figure 4h-j**). In addition to sucrose degradation, these six MAGs covered a range of metabolic pathways, including fixation of CO_2_ and N_2_, the use of reduced sulfur compounds and hydrogen as energy sources, and sulfate respiration (**Table S6; Supporting Information**). Both MAG 76 and 154 are at least facultative anaerobes based on the absence of cytochrome c oxidase in their genome. Given our results that phenolic compounds inhibited microbial sucrose degradation under anaerobic conditions, we predicted that anaerobic sucrose degraders should be able to degrade phenolics. Indeed, expression data indicated that both MAG 76 and 154 degrade phenolics through the benzoyl-CoA and beta-ketoadipate pathways (**Table S6; Supporting Information**). Furthermore, expression data for three of the four aerobic MAGs indicated that these also degrade phenolics, allowing them to metabolize sucrose in the upper layers of the seagrass rhizosphere (see **Supporting Information**).

Our metagenomic and metatranscriptomic results are consistent with our sediment incubation experiments. Many members of the microbial community underneath the meadow have the genomic ability to degrade sucrose. However, low transcription of genes involved in sucrose metabolism indicate most members of the seagrass rhizosphere do not actively degrade this carbon source. We hypothesize that the few taxa that disproportionately expressed genes for sucrose degradation over other sugars evolved mechanisms that enabled them to thrive on this abundantly available and easily degradable energy and carbon resource in the seagrass rhizosphere in spite of the presence of phenolic compounds. This follows the wealth of data from terrestrial rhizospheres showing that select microorganisms are able to use sugars and other compounds in the presence of phenolics^43, 44, 48^. In fact, in terrestrial rhizospheres, phenolic compounds work to shape soil community composition by acting as antimicrobials towards some microbial taxa while stimulating others^48–50^.

### Sucrose exudates are equal to 25% of *P. oceanica’s* net ecosystem metabolism

To contextualize our findings within the larger framework of *P. oceanica* carbon cycling, we compared respiration rates of sucrose measured in this study to the net ecosystem metabolism (photosynthesis minus respiration) of a Mediterranean *P. oceanica* meadow off Elba, Italy during high productivity^29^. We used the potential respiration rates from our sediment incubations to estimate how quickly the calculated standing stock of sucrose of 4.8 mmol/m^2^ in the upper 30 cm underneath a *P. oceanica* meadow would be completely respired to CO_2_ in the absence and presence of seagrass-derived phenolics (**for calculations see Table S3**). In the absence of phenolic compounds, the sucrose pool underneath the meadow would be completely respired to CO2 at areal rates between 64 (anoxic conditions) and 232 (oxic conditions) mmol(C) m^-2^d^-1^, resulting in the complete removal of sucrose to CO2 in less than one day (**Table S3**). Considering previous measurements of *P. oceanica’s* gross primary production at 162 mmol(C) m^-2^d^-1^ (GPP) and net ecosystem metabolism of 77.1 mmol(C) m^-2^d^-1^ (NEM)^29^, the meadow’s productivity would not be able to support these sucrose respiration rates. In fact, the potential rates under anoxia, which represents the majority of the rhizosphere, would be over 205% of the plants daily NEM. These data support our hypothesis that to maintain the high sucrose pool observed underneath the meadow, there must be a strong metabolic inhibitor of microbial respiration. Indeed, our incubation results showed the presence of seagrass-derived phenolics under anoxia led to an approximate 10-fold reduction in the consumption of sucrose and consequent reduced production of CO2 under anoxic conditions, with carbon respired at a rate of 20 mmol(C) m^-2^d^-1^ (**Figure 3c**). At this rate of sucrose removal, which is likely the dominant rate in the largely anoxic, phenolic-rich rhizosphere, we estimate sucrose respiration to account for 25.9% of the total respiration driven by *P. oceanica* primary production. Because the amount of sucrose respired represents the minimum amount that must be excreted by the plant to maintain the high sucrose concentrations in the rhizosphere, we conservatively estimate *P. oceanica’s* release of sucrose to the sediments makes up one quarter of the seagrass meadow NEM (77.1 mmol(C) m^-2^d^-1^)(**Figure 5, Supporting Information**).

**Figure 5.**
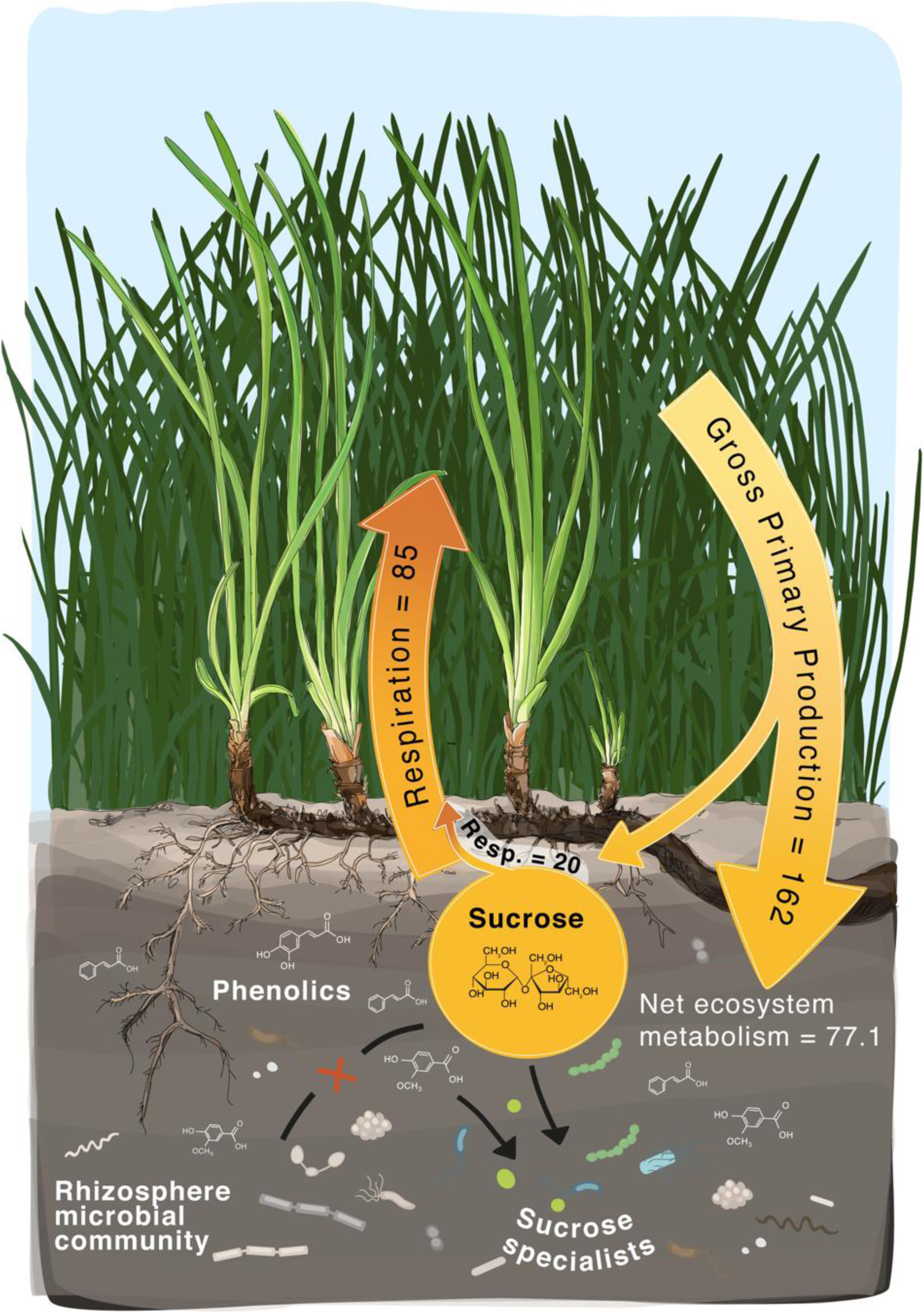
Sucrose release represents over 25% of *P. oceanica’s* net ecosystem metabolism. An overview of ecosystem metabolism underneath a *P. oceanica* meadow. Our combined metabolomics, metagenomics, metatranscriptomics, and incubation results showed the presence of phenolics limits most microbial respiration of sucrose with the exception of select taxa that express genes for degrading plant-produced phenols. Using previously published gross primary production, respiration, and ecosystem metabolism rates^29^, we show our sediment incubation rates from the anoxic, phenolic incubations represent 25.6% of the net ecosystem metabolism of a *P. oceanica* meadow. Arrows representing respiration, gross primary production, and respiration from sucrose are proportional in area to measured rates (mmol(C) m^-2^ d^-1^).

## Conclusions

Our study revealed that a considerable amount of organic carbon derived from seagrasses in the form of sucrose does not enter the microbial loop, which likely has significant consequences for carbon burial in these critical coastal ecosystems. Given that a large portion of seagrass primary production is deposited into sediments daily as a reactive carbon source, we predict that if intact meadows are destroyed, or significant sediment oxygenation events occur, this carbon would be quickly consumed and released to the atmosphere as CO_2_. On a global basis, if seagrasses were completely removed from coastal ecosystems, the oxidation of sucrose alone would release between 0.21 and 0.42 Tg of carbon producing between 0.77 and 1.54 Tg of CO_2_. These results provide evidence that the reactivity of dissolved organic matter is complex and governed by multiple biotic and abiotic processes, including the metabolic potential of the microbial community, the availability of electron acceptors, and inhibitors of microbial metabolism.

As a valuable energy resource, why do seagrasses release sucrose to their sediments? Seagrass release of sugars is most likely an evolutionary mechanism to deal with overflow metabolism during periods of high productivity. Seagrasses evolved a reduced capacity to produce structural carbohydrates, such as cellulose, in favor of maintaining a large osmolyte pool of nonstructural carbohydrates, such as sugars, to compensate for high salinity compared to their terrestrial relatives^51^. Seagrass release of excess photosynthate to their sediments in the form of sucrose could also serve as an overflow valve when productive rates exceed storage capacity. In most marine anoxic environments, plant release of sugars would stimulate sulfate reduction by sediment microbes leading to the accumulation of high concentrations of sulfide, which is toxic to seagrass roots^52–54^. However, we show that the presence of phenolics limits the microbial respiration of sugars under anoxic conditions. This would have the selective advantage that seagrasses could deposit their excess photosynthates to their sediments, while limiting the formation of sulfide.

Another selective advantage for seagrasses in releasing sucrose is that it could attract key microbial partners that benefit plant fitness within the seagrass rhizosphere. In terrestrial plants, during their early developmental stages sucrose makes up 50% of their exudates, and it has been suggested this early priming of the plant’s rhizosphere with sugar is critical in shaping plant-associated microbial communities^42^. Plant exudation of sugars and other compounds is tied to chemotaxis of potential symbionts, including soil bacteria and mycorrhizal fungi^48^. In our study, we show that in the seagrass rhizosphere, the vast majority of sediment microorganisms are not able to effectively metabolize sucrose. However, we did identify a select set of sucrose specialists, that under high sucrose and phenolics concentrations, can both breakdown phenolics and consume sucrose. Some of these sucrose specialists are also capable of sulfur and hydrogen oxidation, and, intriguingly, nitrogen fixation. In oligotrophic waters, such as the Mediterranean, nitrogen is a limited resource for growth. Sucrose release by seagrasses, including *P. oceanica*, to their rhizosphere could support beneficial symbionts that are metabolically active and fix nitrogen in the presence of phenolics and absence of oxygen.

Seagrasses are disappearing at a rate of 110 km^2^ yr^-1^, largely due to anthropogenic activities including land use changes that lead to eutrophication and habitat loss^55^. We predict these harmful activities will disrupt the delicate balance between seagrass excretion and microbial respiration processes. Consequently, it is critical to consider how changes in environmental conditions alter plant-microbial interactions in marine rhizospheres, and the capacity of seagrasses to sequester carbon.

## Methods

### Porewater sampling

Porewater was collected from vegetated and non-vegetated sediments for metabolomic analyses from *P. oceanica* seagrass meadows in the Mediterranean off the island of Elba, Italy (42° 48’29.4588” N; 10° 8’ 34.4436” E), Caribbean at Carrie Bow Cay, Belize (N 16° 04’ 59”; W 88° 04’ 55”) and Twin Cayes, Belize (N 16° 50’ 3”; W 88° 6’ 23”), and Baltic Sea off the coast of Kiel, Germany (54° 27’ 26.56256” N; 10° 11’ 33.1908” E). Using a steel lance (1 m long, 2 µm inner diameter) outfitted with a wire mesh (63 µm) to prevent the intake of sediment and seagrass, porewater was slowly extracted from sediments into sterile syringes. A porewater profile consisted of top to bottom sampling of the sediments every 5 or 10 cm down to 30 or 40 cm depending on sampling location. Following porewater extraction, samples were frozen at -20°C until processing for metabolomic analyses.

Porewater profiles were collected at higher spatial and temporal scales from two *P. oceanica* meadows off the island of Elba, Italy. Profiles (n=3) from replicate (n=3) transects were collected inside, 1 m at the edge and 20 m outside a meadow in Sant’Andrea Bay, Italy (42° 48’29.4588” N; 10° 8’ 34.4436” E ; 6-8 m water depth). These collections were repeated in April, July, and October of 2016 to describe metabolite composition across *P. oceanica’s* growing season.

To explore how porewater metabolite composition changes across a diurnal cycle, porewater samples were collected at 10 cm sediment depth during a 24 hour diurnal cycle in Galanzana Bay, Elba, Italy (42° 44’9.438” N; 10° 14’ 16.3032” E; approximately 2 m water depth). Using replicate lances fixed in the sediment throughout the experiment (n=10) sediment water was extracted (2 mL) at time points around high and low solar irradiance periods (10:00, 12:00, 14:00, 20:00, 22:00, 0:00, 2:00 and 5:00) on 23-24 May 2017. Following collection, samples were immersed in liquid nitrogen and subsequently stored at -20 °C until further processing.

### Seagrass sampling

Individual *P. oceanica* plants from Galanzana Bay were sampled during a diurnal cycle (at 10:00, 12:00, 14:00, 20:00, 0:00, and 5:00) in parallel with sediment porewater samples. Immediately following collection (< 5 min), whole plants were immersed in liquid nitrogen to halt changes in metabolite composition. Plants were stored at -80 °C until extraction.

### Gas chromatography-mass spectrometry based metabolomics

Metabolomic profiles were obtained from sediment porewaters and prepared for gas chromatography-mass spectrometry (GC-MS) using a recently described method^56^.

Metabolites were extracted from seagrass tissues using a modified method for plant-based metabolite profiling^57^. Briefly, tissues from frozen plants were separated into leaves, rhizomes, and roots and ground under liquid nitrogen to a fine powder using mortar and pestle. Approximately 400 mg of ground plant tissue was aliquoted into Eppendorf vials. Pre-cooled methanol (1.4 mL) was added to each aliquot and vortexed for 10 s. Tissues were extracted at 70 °C for 10 min in a thermomixer at 950 rpm and subsequently centrifuged to pellet the tissue at 11,000 *g* at 4 °C. The supernatant was transferred to 14 mL glass vials. To separate the polar and non-polar compounds, 750 µL of chloroform and 1.5 mL of high-grade water was added to each sample, vortexed for 10 s and centrifuged for 15 min at 2,200 *g*. 25 µL of the upper, polar phase was dried to completeness in a vacuum concentrator without heating (approximately 1 h) after which the aliquots were stored at -80 °C until metabolite derivatization.

To remove condensation formed during extract storage, directly before preparation for GC-MS analysis we further dried extracts in a vacuum concentrator for 30 min. Metabolite derivatization was performed by adding 80 µL of methoxyamine hydrochloride dissolved in pyridine (20 mg mL^-1^) to the dried pellet and incubating for 90 min at 37 °C using a thermal rotating incubator under constant rotation at 1350 rpm. Following recent advancements in signal improvement^58^, the pyridine was removed from the sample at room temperature under a gentle flow of N_2_ gas (approximately 1 hour). Following the addition of 100 µL of N,O-B is(trimethylsilyl)trifluoroacetamide, each extract was vortexed, and incubated for another 30 min at 37 °C using a thermal rotating incubator under constant rotation at 1350 rpm. 100 µL was transferred to a GC-MS vial for further GC-MS data acquisition.

#### GC-MS data acquisition

To obtain metabolite profiles from both porewater and seagrass extracts, all derivatized samples were analyzed on an Agilent 7890B GC coupled to an Agilent 5977A single quadrupole mass selective detector. The injector temperature was set at 290 °C. Using an Agilent 7693 autosampler, 1 µL was injected in splitless mode through a GC inlet liner (ultra inert, splitless, single taper, glass wool, Agilent) onto a DB-5MS column (30 m × 0.25 mm, film thickness 0.25 µm; including 10 m DuraGuard column, Agilent). The inlet liner was changed after every 50 samples when processing porewater extracts and after 100 samples when processing seagrass tissues to avoid damage to the GC column. Metabolite separation on the column was achieved with an initial oven temperature of 60 °C followed by a ramp of 20 °C min^-1^ until 325 °C was reached, which was then held for 2 min. Helium carrier gas was used at a constant flow rate of 1 mL min^-1^. Mass spectra were acquired in electron ionization mode at 70 eV across the mass range of 50–600 *m/z* and a scan rate of 2 scans s^-1^. The retention time was locked using standard mixture of fatty acid methyl esters (Sigma Aldrich).

#### Metabolome data analysis

To identify differences in metabolites between vegetated and non-vegetated sediments, we used a non-targeted metabolomic approach to assess porewater samples collected inside and outside a *P. oceanica* seagrass meadow. Raw GC-MS data from samples collected in October 2016 were imported into R (v3.5.2) as converted mzXML files (MsConvert)^59^ and processed using XCMS (v2.99.6)^60^. Individual peaks were picked using the matchedFilter algorithm with a full width at half maximum set to 8.4, signal to noise threshold at 1, *m*/*z* width of 0.25 (step parameter), and *m*/*z* difference between overlapping peaks at 1. Resulting peaks were grouped, retention times corrected and regrouped using the density (bandwidth parameter set to 2) and obiwarp methods. Following peak filling, the CAMERA (v1.32.0)^61^ package was used to place *m*/*z* peaks into pseudo-spectra by grouping similar peaks with the groupFWHM function. A single *m/z* value was selected to represent each CAMERA group using the following criteria: 1) *m/z* value > 150, 2) occurs across samples with the highest frequency and, if two or more ions are tied in terms of number of samples detected, 3) has the highest mean intensity. Ions with very low mean intensities (< 0.001) were considered noise and removed from the analysis. Ion abundances were normalized to the ribitol internal standard and compared using a volcano plot to show ions with significant differences (alpha < 0.1) between sampling location and high fold change values (log(FC) > 2). These peaks were subsequently identified using the Mass Hunter Suite and through comparison to the NIST database.

Using our non-targeted analysis as a guide, identified sugars were further quantified in both porewater and seagrass metabolomic profiles using Mass Hunter Quantification Suite (Agilent). Salinity matched calibration curves were used to determine absolute sugar concentrations in porewater samples. For all statistical analyses, calculated sugar concentrations above the instrument detection limits were conservatively assigned the maximum calibration concentration (200 µM). Values below the detection limits were considered noise and given a value of 0 µM. The relative ion abundances of plant sugars were normalized to tissue wet weight and compared using an analysis of variance (ANOVA) across sampling time. Sugar concentrations from sediments were compared across spatial and temporal scales using ANOVA models where sugar concentration was a function of (1) sampling location*sampling depth, (2) sampling month and (3) time of day blocked by sampling spot. Sugar concentrations were transformed to meet the assumptions of the model.

### Estimation of global sucrose abundances

To determine the amount of sucrose underneath seagrass meadows, the minimum and maximum concentrations across all sampling depths underneath each of the four types of meadows investigated were used to provide a conservative range. These values were extrapolated to the known area of seagrass meadows worldwide^1^. By calculating the total L of porewater given a porosity of 0.42 within the upper 30 cm underneath seagrass meadows, we estimated the total g of sucrose occurring below seagrasses on a global scale using the following equation:

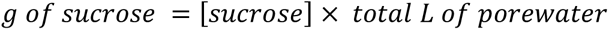

### DOC sampling & analysis

Dissolved organic carbon (DOC) samples were collected in parallel with porewater metabolomic samples from inside, at the edge and 20 m outside a *P. oceanica* seagrass meadow in October of 2016. Taking care to avoid oversampling porewater space, approximately 20 mL of each porewater sample was collected into polypropylene syringes. Samples were filtered through pre-combusted (500 °C, 4 h) Whatman GF/F filters (0.7 µm) into 20 mL acid-washed and pre-combusted scintillation vials. Samples were acidified to pH 2 using 25% hydrochloric acid and stored at 4 °C until analysis. DOC concentrations were measured by high temperature catalytic oxidation on a Shimadzu TOC-VCPH analyzer. Instrument trueness and precision was tested against Deep Atlantic seawater reference material (42 µM DOC; acquired from the Hansell lab at University of Miami, FL, USA) and low carbon water (1 µM DOC) and was better than 5%.

### Molecular DOM analysis

DOM was extracted from filtered (0.7 µm GF/F) and acidified (pH2) porewater samples using solid-phase extraction^62^. Briefly, Agilent Bond Elut PPL (0.1 g) cartridges were conditioned with methanol followed by ultrapure water (pH 2 using 25% hydrochloric acid). 15 to 20 mL of acidified porewater (pH 2) was loaded onto the cartridge followed by another rinse with ultrapure water (pH 2). The cartridges were dried with N_2_ gas and DOM was eluted with 0.5 mL methanol and stored at −20 °C until further analysis. The extraction efficiencies were determined as DOC in methanol extract divided by bulk DOC concentrations of the original porewater samples. DOC in methanol extracts was determined by drying an aliquot of extract and re-dissolving in ultrapure water. The extraction efficiency was 37 ± 11%. Methanol extracts with isolated DOM were diluted in 1:1 methanol/water to a final concentration of 5 mg C L^−1^. The samples were analyzed by ultrahigh-resolution mass spectrometry as previously described^63^ using a solariX XR Fourier transform ion cyclotron resonance mass spectrometer (FT-ICR-MS, Bruker Daltonik GmbH, Bremen, Germany) connected to a 15 T superconducting magnet. The samples were analyzed in negative mode and infused into the electrospray source (Apollo II ion source, Bruker Daltonik GmbH, Bremen, Germany) at 2 μL min^−1^. Two hundred scans were collected with a mass window from 92 to 2000 Da. Molecular formulae above the method detection limit^64^ were assigned with the following restrictions: ^12^C_1−30_ ^1^H_1−200_ O_1−50_ ^14^N_0−4_ S_0−2_ P_0−1_. Molecular masses were removed for further analysis when detected in less than three samples. Samples were normalized to the sum of FT-ICR-MS signal intensities and a modified aromaticity index (AI_mod_)^65^ was calculated for each sample. Molecular formulae were categorized as highly aromatic if 0.66 ≥ AI_mod_ ≥ 0.50 which includes molecular formulae with element ratios that are consistent with polyphenols^63^.

### Seagrass root mass spectrometry imaging

Seagrass root pieces were embedded into pre-cooled (4 °C) 2% carboxymethylcellulose blocks and snap frozen in liquid nitrogen. Roots were cross-sectioned into 12 µm thick slices with a cryrostat (Leica CM3050 S, Leica Biosystems Nussloch GmbH; chamber and object holder temperature at -25 °C). Individual sections were obtained by pressing the sample between two frozen Polylsine glass slides to ensure a flat surface for mass spectrometry imaging (MSI) analysis. Before MSI sample preparation, overview images of the root sections were acquired with a stereomicroscope (SMZ25, Nikon, Düssedorf, Germany) and, for the high-resolution figure in the manuscript, with an Olympus BX53 compound microscope (Olympus, Tokyo, Japan) at a resolution of 0.36 µm/ pixel, operated through the software cellSens (Olympus, Tokyo, Japan). For matrix application, super 2,5-dihydroxybenzoic acid solution (S-DBH; 30 mg mL^-1^ in 50% methanol:water with 0.1% trifluoroacetic acid (TFA) was applied to each section using an automated pneumatic sprayer (SunCollect, SunChrome, Wissenschaftliche Geräte GmbH, Germany). Spraying consisted of 10 layers at a flow rate of 10 µL min^-1^ (layer 1) and 15 µL min^-1^ (layers 2-10) resulting in a homogeneous layer of crystalline matrix with crystal sizes around 1 µm. To identify phenolic compounds within seagrass root tissues, an additional seagrass root section was prepared by applying 9-aminoacridine matrix (9AA) in 70% methanol:water with 0.1% TFA using an automated pneumatic sprayer. Spraying consisted of 8 layers in total at a flow rate of 10 µL min^-1^ (layer 1), 20 µL min^-1^ (layer 2), 30 µL min^-1^ (layer 3) and 40 µL min^-1^ (layers 4-8).

MS-imaging was done with an atmospheric pressure matrix-assisted laser desorption/ionization (AP-MALDI) ion source (“AP-SMALDI10”, TransMIT GmbH, Gießen, Germany), coupled to a Fourier transform orbital trapping mass spectrometer (Q Exactive^TM^ HF, Thermo Fisher Scientific GmbH, Bremen, Germany). MS images were collected by scanning the matrix-covered tissue sections at a step size of 10 µm without overlapping of the laser spots. Mass spectra were acquired in positive-ion mode for all S-DBH prepared sections with a detection range of *m/z* 50 to 750 and a mass resolving power of 240,000 at *m/z* 200. Sections prepared with 9AA matrix were acquired in negative-ion mode with a *m/z* range of 70 to 1000 at a 20 µm step size.

### Oxygen detection

Oxygen profiling within the seagrass bed was carried out in situ using a combination of custom built and commercially available instruments. Briefly, an oxygen flow-through cell (Pyroscience, OXFTC) was connected to a stainless steel push-point sampling lancet. The inside of the oxygen flow-through cell is coated with an oxygen-sensitive fluorescent dye, which allows contactless oxygen measurements from outside of the cell. Dye excitation and emission readout was performed via a custom-made glass fiber connected to an underwater oxygen OEM module (Pyroscience, FSO2-SUBPORT). The oxygen module data was stored using an analog logger from a diver-operated motorized microsensor profiler system (DOMS, see^66^). Prior to sampling, the oxygen flow-through cell was calibrated using a two-point calibration procedure. First, aerated sea water was percolated through the chamber (100%) followed by sea water treated with natrium dithionate (0%). Scuba divers collected two porewater profiles down to a depth of 30 cm in 2.5 cm intervals. In contrast to previous studies^67^, our method did not allow us to detect microscale gradients in subsurface peaks in oxygen close to seagrass roots.

### Microbial activity

Sediments were collected using cores inside (n=3) and at the edge (n=3) of a *P. oceanica* meadow in Sant’Andrea Bay, Elba, Italy (42° 48’29.4588” N; 10° 8’ 34.4436” E ; 6-8 m water depth). Individual cores were subsampled inside a glove bag under nitrogen gas to avoid introducing oxygen to the anoxic sediment layers. Cores were split into oxic (upper 3 cm) and anoxic fractions (between 5 and 10 cm). Individual fractions were mixed and 10 mL of sediment volume were added to each serum vial. Replicate 250 mL serum vials for each fraction were filled with filtered seawater (environmental control), filtered artificial seawater (technical control), or filtered artificial seawater with approximately 35 µM of phenolic extract prepared from seagrass tissue as previously described^68^. We introduced an equal amount of phenolics and sugars as this ratio was reported for other seagrass leachates^41^. Assuming the main phenolic compound is chicoric acid^69^, this corresponds to a concentration of approximately 35 µM. All serum vials had 100 mL of headspace with either N_2_ (anoxic) or air (oxic). To remove background levels of phenolics and other compounds, sediments incubated with artificial seawater were flushed with five times the sediment volume in artificial seawater before transfer to serum vials. Fully labeled ^13^C_12_-sucrose was added to each bottle at a final concentration of 50 µM. Oxic incubations were prepared in ambient air, with optode spots confirming the presence of oxygen throughout the experiment. Anoxic incubations under nitrogen atmosphere and both natural and artificial seawater were bubbled with N_2_ gas until < 2 µM of oxygen was measured.

Incubation seawater from each serum bottle was collected for GC-MS based metabolomics and cavity ring-down spectrometer (CRDS). At each time point, 5 mL of incubation seawater was collected from each vial and replaced by either filtered air (oxic) or N_2_ gas (anoxic). 2 mL was transferred to an Eppendorf tube and stored at -20 °C until analysis (GC-MS), while the remaining 3 mL were transferred to a nitrogen filled exetainer with 50 µL of saturated mercury(II) chloride to halt microbial metabolism. All samples were evidence indicated microbial metabolism was not successfully halted (cross validated using metabolomics results) were excluded from the CRDS analysis.

The metabolic activity of the sediment was assessed by monitoring the disappearance of ^13^C_12_-sucrose (GC-MS) and the production of ^13^CO_2_ (CRDS) over time. Samples were prepared for GC-MS following the metabolomics protocol for seawater described above. To determine the rate of ^13^CO_2_ production from ^13^C_12_-sucrose, 3 mL of each sample was acidified with 50 µl 20% phosphorphic acid^70^ in preparation for cavity ring-down spectroscopy (G2201-i coupled to a Liaison A0301, Picarro Inc., connected to an AutoMate Prep Device, Bushnell) . The production of ^13^CO_2_ from ^13^C-sucrose was calculated by subtracting the initial ^13^CO_2_ concentration from each subsequent time point for each bottle. The production of ^13^CO_2_ from ^13^C_12_-sucrose was calculated from the increase of ^13^CO_2_ (in ppm) over total CO_2_ (in ppm) over 24 hours. The initial ratio of ^13^CO_2_/^12^CO_2_ was then subtracted from the subsequent time points and the difference was multiplied by the CO_2_ concentration in the sample.

Volumetric sucrose oxidation rates were determined by linear regression of the measured ^13^C_12_-sucrose and ^13^CO_2_ concentrations. The rates from the incubations were corrected for dilution and sediment porosity assumed as 0.42 (based on previous calculations). A correction based on the measured ^12^C_12_-sucrose concentration was applied to account for dilution of the labeled substrate. For rates per area an active layer of 20 cm thickness was assumed based on the average sucrose concentrations in the sediment and multiplied with the volumetric rates.

### Sediment nucleic acid extraction, sequencing and analysis

Genomic DNA and total RNA were extracted from sediment cores (n=3) collected inside, at the edge, and 20 m outside a *P. oceanica* meadow in Sant’Andrea Bay, Elba, Italy in October 2016. Directly after collection, cores were sectioned into 5 cm slices and frozen at -20 °C. A subsample of each sediment slice was also presevered in RNAlater (SigmaAldrich) for RNA extraction. Samples were stored at -80 °C until further processing.

#### DNA extraction

DNA was extracted from 0.2 g of the 10-15 cm slice. Prior to extraction, extracellular DNA (exDNA) was removed from the sediments following previously described methods^71, 72^. Briefly, exDNA was removed by suspending frozen sediments into 200 µL of carbonate dissolution buffer. The solution was incubated at room temperature for 1 h at 600 rpm, 1.6 mL of 10X tris hydrochloride buffer (300 mM; pH 10) added to each sample, and incubated for an additional hour. All samples were centrifuged at 10,000 x g for 20 min to separate dissolved exDNA from sediments containing intact microbial cells. Remaining cellular DNA was extracted from pelleted sediments in 500 µL of extraction buffer^72^. Following 3 freeze thaw cycles, 10 µL of proteinase K (20 mg µL^-1^) was added to each sample and incubated for 30 min at 37 °C. Each sample was then incubated (2 h, 50 °C) with 50 µL of 20% SDS buffer and then centrifuged at 4000 x *g* for 10 min. After re-extracting the pellet in both buffers (166 µL of extraction buffer and 16 µL of 20% SDS at 65 °C for 20 min), sample supernatants were combined with equal volumes of chloroform: isoamylalcohol (24:1 v/v) and centrifuged (10,000 x *g*, 10 min). This step was repeated a second time and the resulting aqueous phase containing DNA was combined with 600 µL of isopropanol and stored overnight at 4 °C. To pellet DNA, samples were centrifuged (14,000 x *g*, 30 min) and the supernatant discarded. The DNA pellet was washed in pre-cooled 80% ethanol and centrifuged (14,000 x *g*, 10 min). The resulting supernatant was removed and the DNA pellet was dried in a vacuum concentrator (Eppendorf concentrator plus, 10 min, V-AL). DNA was eluted in DEPC water and stored at -20 °C until metagenomic library preparation.

#### RNA extraction

Total RNA was extract from a separate aliquot of the 10-15 cm slice of each sediment core preserved in RNAlater following a modified version of previously described methods for RNA recovery from marine sediments^73^. Briefly, 0.2 g of each sediment sample was homogenized in 500 µL of lysis solution I (30 mM Tris-HCL, 30 mM EDTA, 800 mM guanidium hydrochloride, 0.5% TritonX-100) and flash frozen in liquid nitrogen. Samples were thawed at 65 °C under gentle agitation, and then vortexed for 10 s and incubated for 60 min at 50 °C under constant shaking. The resulting mixture was flash frozen in liquid nitrogen and thawed at 65 °C, after which 500 µL of lysis solution II (2.5 M NaCl, 2% CTAB, 0.1% PVPP) was added to the mixture. Following two more freeze-thaw cycles, samples were incubated for 60 min at 65 °C. Samples were centrifuged for 30 min 5242 x *g* at 4 °C and resulting supernatant transferred for purification. Supernatants containing the nucleic acids were purified using two cycles of washing in 1 volume of chloroform:isoamylalcohol (24:1), inverting 60 times, and centrifuging the mixture for 30 min at 5242 x *g* at 4 °C. The nucleic acids were precipitated from the resulting aqueous supernatant using the following steps: fully dissolve linear polyacrylamide was added to each sample to achieve a final concentration of 20 µL mL^-1^, samples were homogenize with 0.1 volumes of 5 M sodium chloride solution following by 1.5 volumes of isopropanol, this mixture was incubated in the dark at -20 °C overnight, and centrifuged for 30 min at 14,000 x *g* at 4 °C. The nucleic acids were washed by adding 70% ethanol and centrifuged for 10 min at 14,000 x *g* at 4 °C. The pellet was air dried and dissolved in 100 µL of RNAase free water. Following this purification step, the resulting RNA was further cleaned using the Norgen Biotec CallAll kit (BioCat # 23800-NB) following manufacturer’s protocol.

#### Library preparation and sequencing

Library preparation and sequencing was performed at the Max Planck Genome Center Cologne, Germany (https://mpgc.mpipz.mpg.de/home/). Briefly, 10 ng of genomic DNA was fragmented via sonication (Covaris S2, Covaris) followed by library preparation with NEBNext Ultra II FS DNA Library Prep Kit for Illumina (New England Biolabs). DNA library preparation included 6 cycles of PCR amplification. Total RNA was used for cDNA generation Ovation RNA-Seq system V2 (NuGEN) according to the manufacturer’s specifications. Subsequently, 500 ng cDNA was fragmented via sonication, followed by library preparation with NEBNext Ultra II FS DNA Library Prep Kit for Illumina. Library preparation included 5 cycles of PCR amplification. Quality and quantity were assessed at all steps for all libraries via capillary electrophoresis (TapeStation, Agilent Technologies) and fluorometry (Qubit, Thermo Fisher Scientific). All libraries were immobilized and processed onto a flow cell with cBot (Illumina) and subsequently sequenced on HiSeq3000 system (Illumina) with 2 x 150 bp paired end reads.

#### Full length 16S rRNA

The sediment DNA was purified with DNeasy PowerClean Pro Cleanup Kit, Qiagen. 16S amplicons were amplified with the universal primer pair GM3F (5′-AGAGTTTGATCMTGGC-3′) /GM4R (5′-TACCTTGTTACGACTT-3′)^74^ and Phusion DNA polymerase, following the New England Biolabs routine PCR protocol including the addition of DMSO with annealing temperature for the first two cycles at 42 °C, 20 s followed by 45 °C, 20 s for the remaining 34 cycles. The number of PCR cycles for sufficient DNA for downstream applications was empirically determined by quantitative PCR (ViiA 7 Real-Time PCR System, ThermoFisher) in the presence of 1X Evagreen (20X Evagreen, Biotium, Fremont, USA). Amplicons were purified with DNA Clean & Concentrator Kits, Zymo Research or AMpure Beads (Beckman Coulter) and quantified by Qubit HS kit (ThermoFisher). Next, PacBio libraries were prepared as recommended in protocol "2 kb Template Preparation and Sequencing" of Pacific Biosciences using adaptors with 16mer barcode sequence to demultiplex sequencing data. Libraries were pooled equimolar and sequenced on Sequel I of Pacific Biosciences on a single cell with 20 hours run time, polymerase binding kit 3.0, sequencing chemistry version 3.0 and SMRT cell 1Mv3. Resulting reads were error corrected using circular consensus (CCS) reads (10 passes) using the SMRTlink (v 6.0.0) software.

Resulting 16S CCS reads were processed using Dada2^75^ in R to obtain amplicon sequence variants (ASVs). Briefly, amplicon reads were imported into R and filtered to control for expected sequence length (between 1000 and 1600 bp) and high quality reads (minQ=3). Following error estimation, taxonomy was assigned to each unique ASV by comparison against the GTDB database. The resulting count table was imported into phyloSeq^76^ for subsequent analyses including estimate of sequence richness and comparison of taxonomic profiles across samples.

#### Taxonomic composition

phyloFlash (v2.0)^77^ was used to assemble the small subunit of the rRNA gene from each metagenomic library to assess taxonomic composition. The resulting sequence counts, taxonomically classified at order level through comparison to the SILVA SSU 111 (July 2012)^78^ database, were imported into R (v3.5.2) and processed using the phyloseq package (v1.26.1)^76^ to facilitate analysis and visualization. Data processing included removal of eukaryotic taxa and normalization of count data to relative abundances. A principle coordinates analysis (PCoA) was used to assess differences in taxonomic compositions among sampling sites.

#### Metagenomic bin recovery

Metagenomic libraries were used to recover metagenome assembled genomes (MAGs) from marine sediments. Using the bbtools suite (v38.34, http://jgi.doe.gov/data-and-tools/bb-tools/) reads were cleaned of Illumina adaptors and low quality ends were trimmed before normalization to a maximum *k-mer* depth of 100X (*k-mer* length of 31). Reads with an average *k-mer* depth less than two were considered erroneous and were discarded. SPAdes (v3.12.0)^79^ was used to error correct all remaining reads, which were then fed into MEGAHIT (v1.1.3)^80^ to generate a co-assembly with *k*-mers from 21 to 51 in steps of 10. Sequencing coverage of each contig was calculated by mapping the error corrected reads back to the co-assembly using bbmap (v38.34, http://jgi.doe.gov/data-and-tools/bb-tools/).

The metagenomic co-assembly was binned using three automated binning programs (MetaBAT v0.32.5; concoct v1.0.0 and MaxBin v2.2.6)^81–83^ and subsequently dereplicated and aggregated with DAS Tool (v1.1.1)^84^. Bin quality was assessed using the CheckM lineage-based workflow method (v1.0.7)(**TableS4**)^85^. Bins with completeness ≥ 50%, contamination scores corrected for strain heterogeneity ≤ 10% were considered to be of medium draft quality and used in subsequent analyses. Resulting MAGs were taxonomically classified using the GTDBTk classify work flow (v0.2.1)^86^. MAGs represented only 2.9% of the metagenomic read set, however their taxonomic identities reflected the relative abundance of the sediment communities occurring across habitats (**Table S5; Figure S7**). Open reading frames (ORFs) from each MAG were predicted with Prodigal (v2.6.4)^87^ and annotated using a combination of reference database searchers: dbCAN (v2.0)^88^, diamond blastp (v0.9.25)^89^, Uniprot^90^, interproscan (v.5.36-75)^91^ against both PFam^92^ and Phobius^93^, and EggNOG mapper (v2.0)^94^.

#### Expression analysis

Metatranscriptomic reads were cleaned of Illumina adaptors and low quality ends were trimmed using the bbtools suite (v38.34, http://jgi.doe.gov/data-and-tools/bb-tools/) before error correction using SPAdes (v3.12.0)^79^. Ribosomal RNA was filtered from cleaned reads using SortMeRNA (v2.1b)^95^. Non-rRNA reads were mapped to all bins and counted in Kallisto (v0.46)^96^. Raw read counts were imported into R (v3.5.2) for analysis and visualization. To correct for varying relative abundances in MAGs across samples, transcript counts were normalized to one million per each MAG in each sample (transcripts per million - TPM). TMP values for genes with identical annotation were summed within samples. Total transcript counts per habitat per MAG were used to compare transcription levels inside, at the edge and outside the meadow.

#### Phylogenomic tree construction

Phylogenomic trees were constructed from select MAGs using GToTree (v1.4.14)^97^. Briefly, genomes related to each MAG (based on GTDBtk taxonomy) and at least three outgroups were downloaded from NCBI RefSeq or Genbank repositories. Genes were called using Prodigal^87^ and annotated through HHMER3^98^ comparisons to a set of 74 bacterial single copy genes (SCGs). Genomes were retained in the analysis if they contained at least 45% of all SCGs. Annotated genes were aligned using MUSCLE^99^ and trimmed with TrimAl^100^. IQ-TREE^101^ was used to calculate a maximum-likelihood tree with ultrafast bootstrap support values from the concatenated alignment which was visualized using the interactive tree of life webserver^102^.

## Data and code availability

Sequence data from this study were deposited in the European Nucleotide Archive under accession numbers PRJEB35096 and PRJEB40297 using the data brokerage service from the German Federation for Biological Data (GFBio)^103^, in compliance with the Minimal Information about any (X) Sequence (MIxS) standard^104^. Metabolomics data were deposited in Metabolights (https://www.ebi.ac.uk/metabolights/) under accession numbers MTBLS1570 (https://www.ebi.ac.uk/metabolights/reviewer374e7886-f35c-41b1-bded-c401bc56432b), MTBLS1610 (https://www.ebi.ac.uk/metabolights/reviewer9aae6076-90af-4ac7-9eb7-514ce30a1ae5), MTBLS1579 (https://www.ebi.ac.uk/metabolights/reviewer377b3751-9dc7-4fb1-8ab9-38b52108712f), and MTBLS1746 (https://www.ebi.ac.uk/metabolights/reviewer1aad3302-d8d0-491e-9ae9-34caa9539e0c). All other datasets are available at zenodo.org under doi: 10.5281/zenodo.3843378. All code used for data processing and analysis is available at https://github.com/esogin/sweet_spots_in_the_sea.

## Extended Figures

**Extended Data 1.**
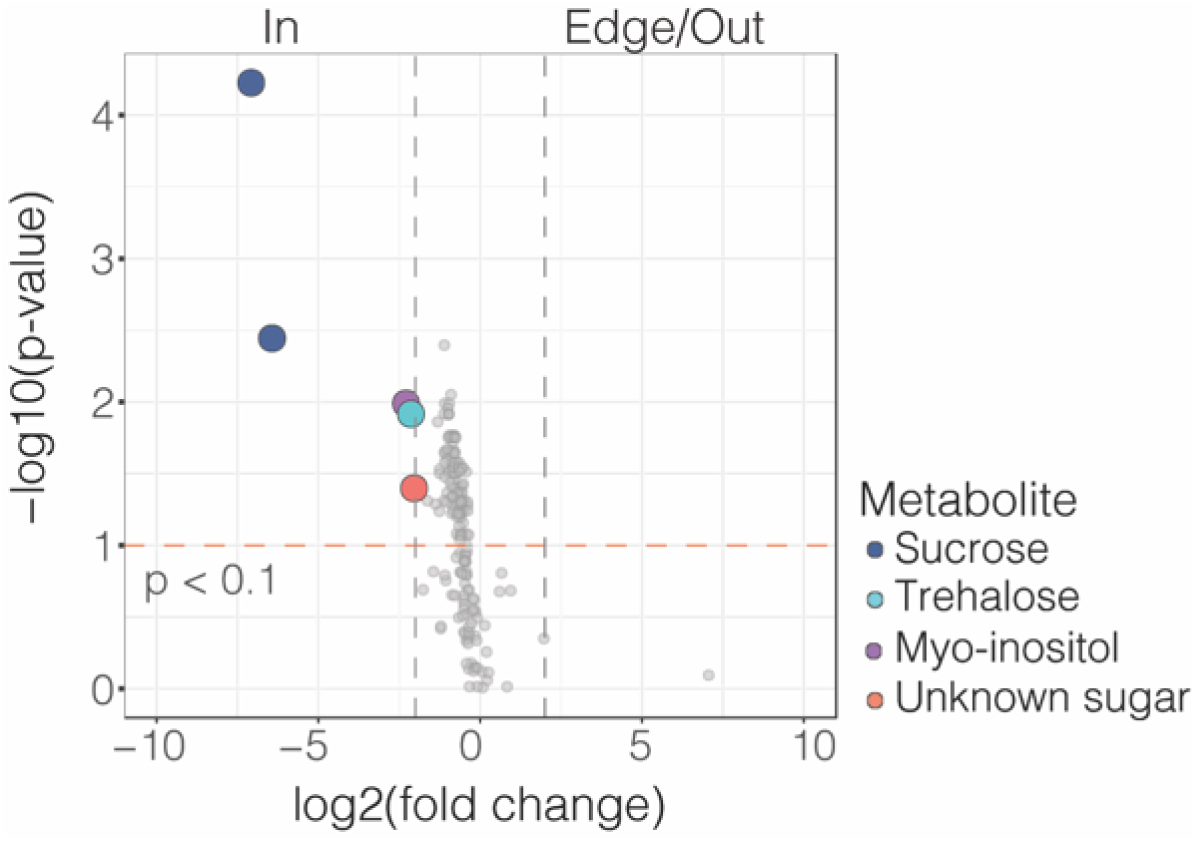
Sugars are more abundant in porewaters underneath seagrass meadows. A volcano plot comparing gas chromatography-mass spectrometry peaks from porewater metabolites collected inside (In) versus 1 and 20 m away from a seagrass meadow (Edge/Out). Significant peaks are represented by colored circles (α < 0.1). The gray dashed lines represent Log2-fold changes > 2; The orange dashed line represents Benjamini-Hochberg corrected p-values < 0.1

**Extended Data 2.**
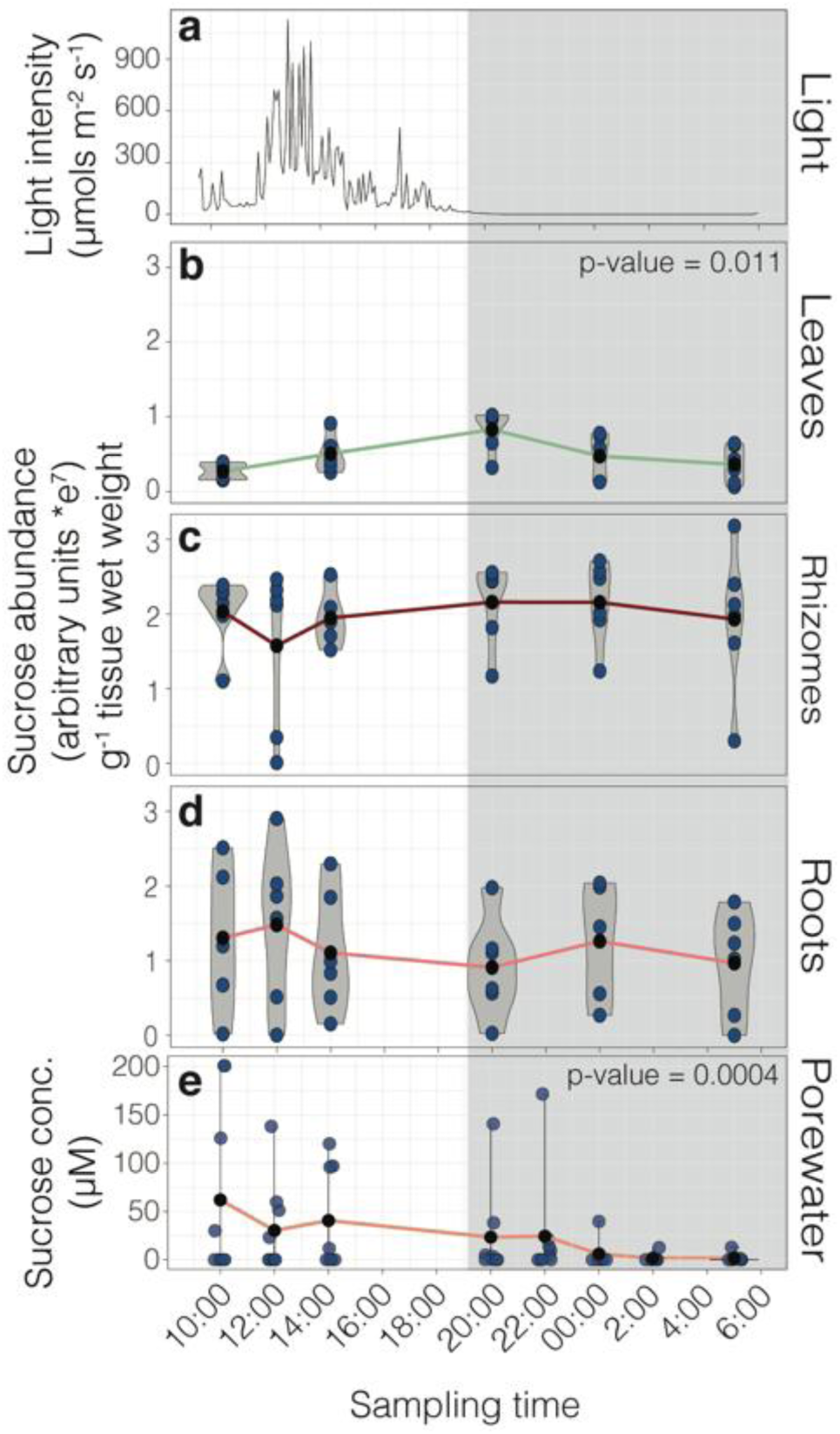
Sucrose abundances vary across a day/night cycle in plant leaves and sediment porewater. **a**, In situ light levels across 24 hours in a Mediterranean *P. oceanica* meadow off the Island of Elba, Italy (> 2 m water depth). **a-d**, Violin plots show the relative abundance of sucrose g^-1^ tissue plant wet weight in **b,** leaves**, c**, rhizomes, and **d,** roots at each sampling time point. Linear models show that sucrose relative abundances were significantly higher in plant leaves at dusk (t= 20:00) than at other times. **e,** Violin plots showing sucrose concentrations measured in sediment porewaters at each sampling time point. A linear mixed effects model with a random effect of sampling spot shows sucrose concentrations significantly changed throughout the day and were higher during the daylight hours then after midnight (t = 0:00) and before dawn (t = 5:00). The means of each sampling time point (black points) are connected by solid lines. For **e,** sucrose concentrations greater than the limits of quantification are represented as 200 µM. All statistical results are reported in **Table S2**. All data points were transformed prior to statistical analyses to meet assumptions of normality.

**Extended Data Figure 3.**
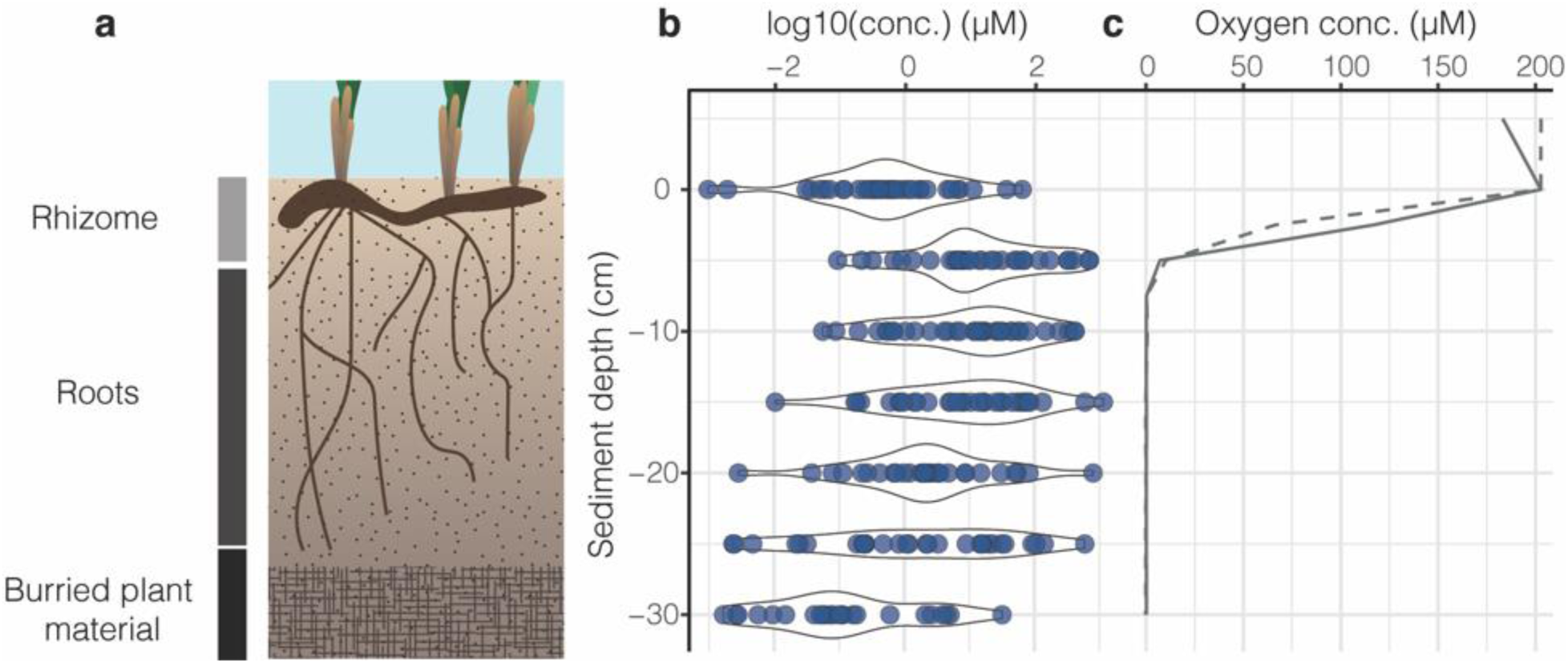
Seagrasses deposit sucrose into their rhizosphere through their roots. a, Schematic showing that *P. oceanica* roots extend between 5 and 20 cm into the sediments in Mediterranean seagrass meadows off the Island of Elba, Italy. **b,** Sucrose concentrations were highest in the zone of root penetration between 5 and 30 cm (*p* = 0.0019, **Table S2**). **c,** Oxygen profiles revealed anoxic conditions below 7.5 cm sediment depth.

**Extended Data Figure 4.**
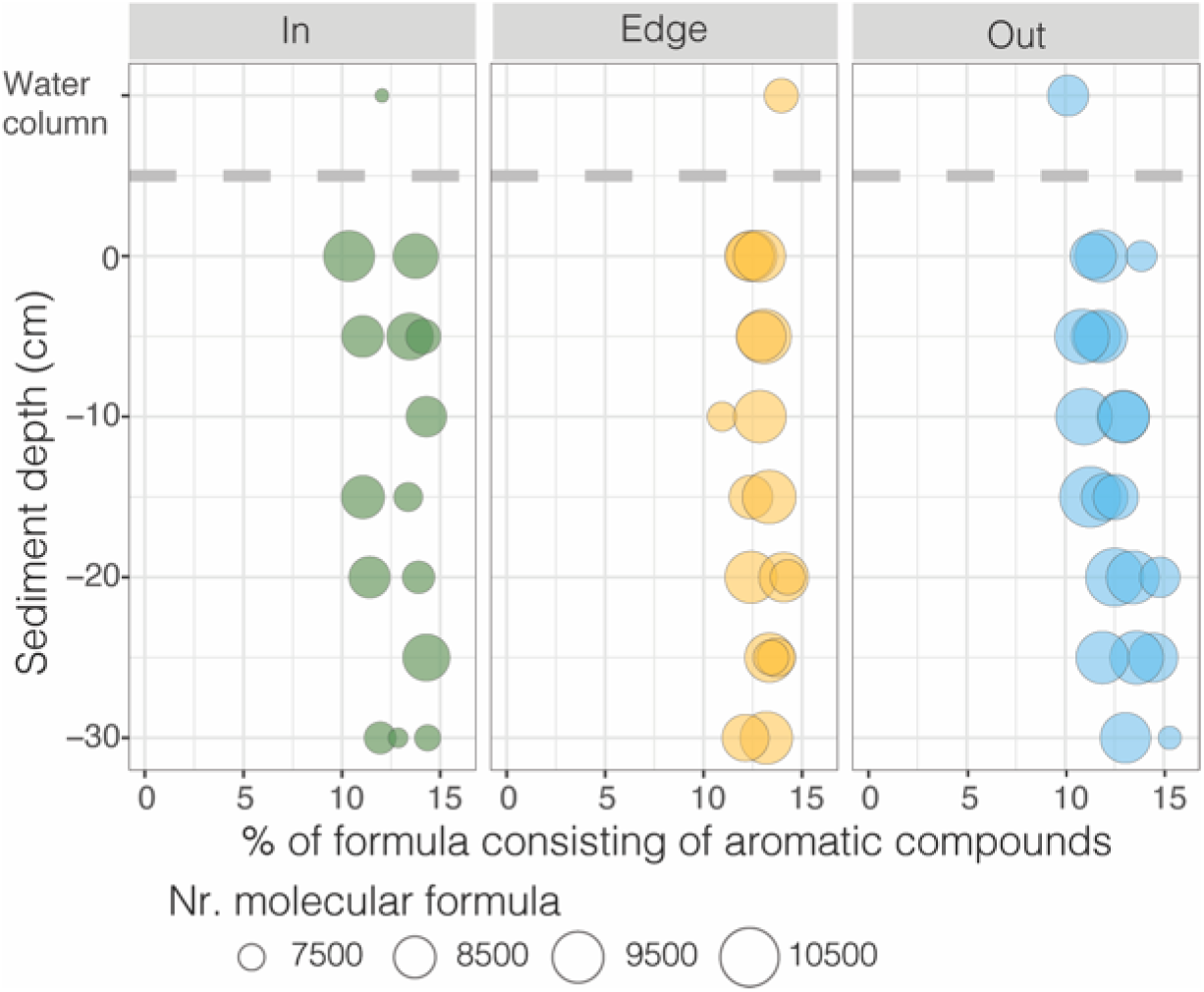
Phenolics are present in sediment porewaters underneath *P. oceanica* meadows. Porewater profiles of the molecular dissolved organic matter composition underneath (In), 1 m (Edge) and 20 m (Out) from seagrass meadows, measured by ultra-high resolution mass spectrometry. Our analyses revealed abundant aromatic molecular formulae, including polyphenols, which comprised between 10 and 16% of all molecular formulae across sampling sites. Size of points reflects number of molecular formula per sample.

**Extended Data Figure 5.**
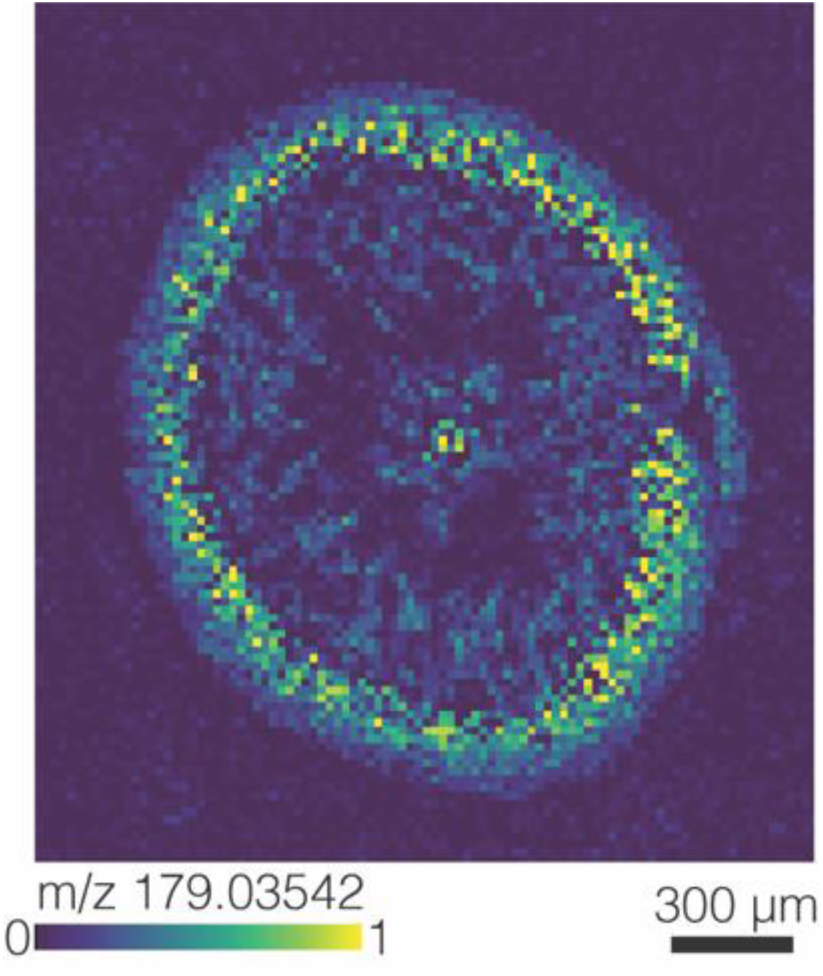
Seagrass roots contain phenolic compounds. Matrix-assisted laser desorption/ionization mass spectrometry image from a Mediterranean *P. oceanica* root cross section collected off the Island of Elba, Italy. The ion image shows the presence of the phenolic compound, putatively identified as caffeic acid (C_9_H_8_O_4_, [M-H]^-^, *m/z* 179.0354). Relative ion abundances are visualized from low (blue) to high (yellow).

**Extended Figure 6.**
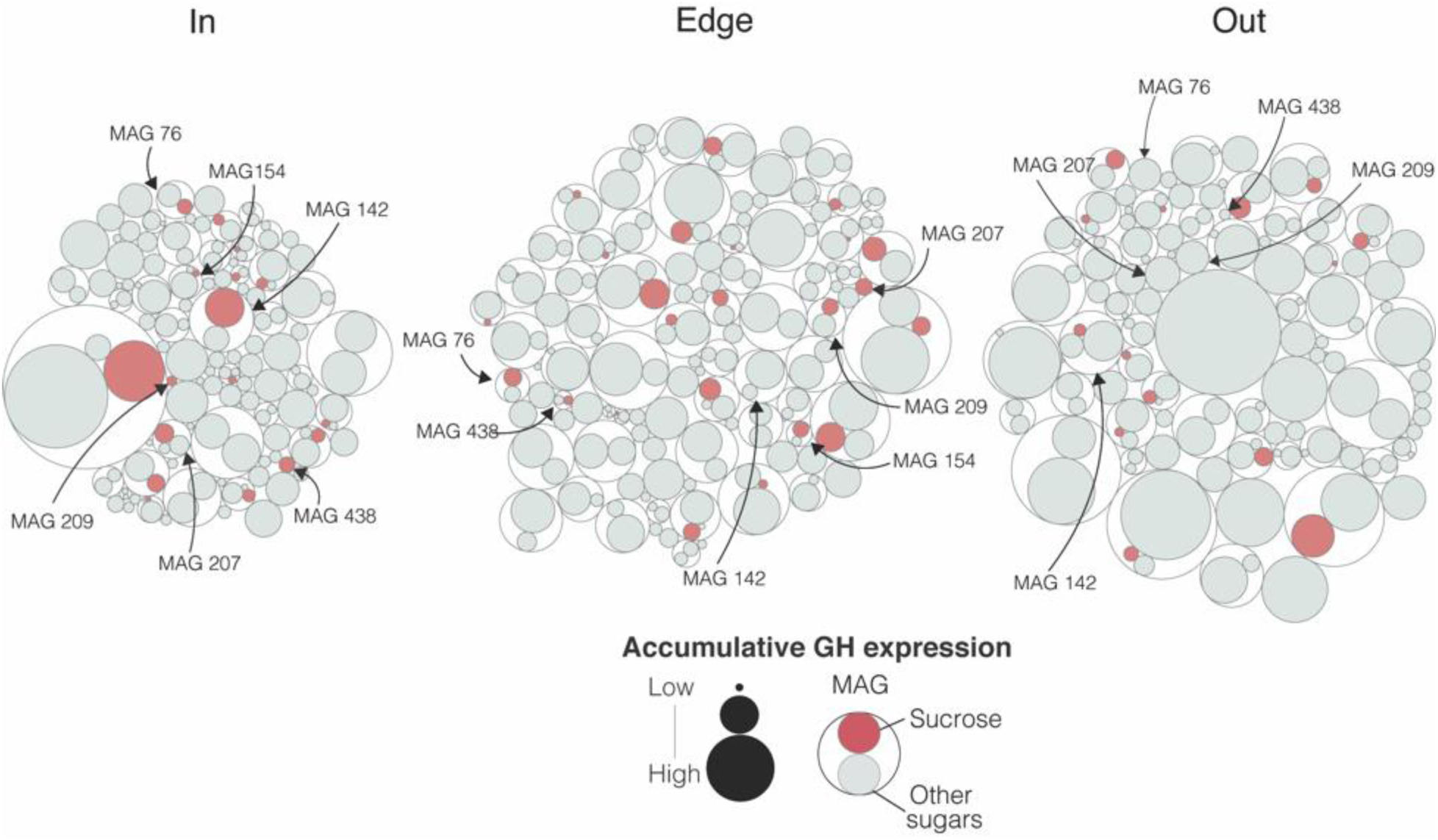
Sucrose specialists show higher expression of genes for metabolizing sucrose than other sugars. Circle packing plots show the hierarchical relationships between the accumulative expression of glycoside hydrolases (GH) for each MAG from underneath (In), 1 m (Edge) and 20 m (Out) away from a *P. oceanica* meadow off Elba, Italy. Inner circles are colored according to the predicted substrates (red = sucrose, light gray = other sugars). The size of the inner circle represents the accumulative transcription level for each MAG (outer white circles) across collection sites. Sucrose specialists are labeled with their MAG numbers.

## Acknowledgements

We thank Tora Gulstad, Kristopher Caspersen, Marvin Weinhold, Frantisek Fojt, Janine Beckmann, Silke Wetzel, and Miriam Sadowski (Max Planck Institute for Marine Microbiology-MPI-MM) for support with data acquisition and sample preparation. We also thank Bruno Huettel (Max Planck Genome Center) for his support with sequencing. We are grateful for fruitful discussions with Marcel Kuypers (MPI-MM), Dirk de Beer (MPI-MM), Dirk Koopmans (MPI-Bremen), Thorsten Dittmar (University Oldenburg), and colleagues in the Departments of Symbiosis and Biogeochemistry (MPI-MM). We thank Miriam Weber, Christian Lott, and HYDRA staff for sample collections. We are grateful to the Max-Planck Society and the Gordon and Betty Moore Foundation (Marine Microbial Initiative Investigator Award to ND, Grant #GBMF3811) for financial support. This work is contribution *XXX* from the Carrie Bow Cay Laboratory, Caribbean Coral Reef Ecosystem Program, National Museum of Natural History, Washington DC.

## Author Contributions

E.M.S., N.D., and M.L. conceived the study. E.M.S. and D.M. collected, processed, and analyzed primary samples for dissolved organic matter, metabolomics, and sequencing analyses. E.M.S preformed statistical analysis for all primary data. M.S. analyzed samples for dissolved organic matter analysis. S.A. collected and analyzed samples for dissolved oxygen concentrations. B.G. collected and analyzed mass spectrometry imaging data. P.B. preformed sediment incubation experiments, and P.B., S.A., and S.S. analyzed data from these incubations. E.M.S., H.G.V. and D.V.M. designed and optimized bioinformatic pipelines for sequencing analysis. D.V.M. and G.D. reconstructed microbial metabolisms based on sequencing results. E.M.S., D.M., H.G.V., N.D., and M.L wrote the initial draft of the manuscript. All authors contributed to review and editing of the manuscript. N.D. obtained funding for the study.

## Notes

### Competing Interest Statement

The authors have declared no competing interest.

### Summary of Updates

This manuscript was revised to update the following: inclusion of transcriptomics data in manuscript, addition of new experimental data and large update to text and figures to clarify central message of the paper.

## References

1. Fourqurean, J. W. et al. Seagrass ecosystems as a globally significant carbon stock. Nat Geoci 5, 505–509 (2012).

2. McLeod, E. et al. A blueprint for blue carbon: toward an improved understanding of the role of vegetated coastal habitats in sequestering CO_2_. Front Ecol Enviro 9, 552–560 (2011).

3. Lange, M. et al. Plant diversity increases soil microbial activity and soil carbon storage. Nat Commun 6, 6707 (2015).

4. Torsvik, V. & Øvreås, L. Microbial diversity and function in soil: from genes to ecosystem. Curr Opin Microbiol 5, 240–245 (2002).

5. Fierer, N. Embracing the unknown: disentangling the complexities of the soil microbiome. Nat Rev Microbiol 15, 579–590 (2017).

6. Fierer, N. & Jackson, R. B. The diversity and biogeography of soil bacterial communities. Proc Natl Acad Sci U S A 103, 626–631(2006).

7. Short, F., Carruthers, T., Dennison, W. & Waycott, M. Global seagrass distribution and diversity: A bioregional model. J Exp Mar Biol Ecol 350, 3–20 (2007).

8. Barbier, E. et al. The value of estuarine and coastal ecosystem services. Ecol Monogr 81, 169–193 (2011).

9. Barrón, C. & Duarte, C. M. Dissolved organic matter release in a *Posidonia oceanica* meadow. MEPS 374, 75–84 (2009).

10. Schmidt, M. W. I. et al. Persistence of soil organic matter as an ecosystem property. Nature 478, 49–56, (2011).

11. Dittmar, T. & Stubbins, A. in Treatise on Geochemistry (Second Edition) (eds Heinrich D. Holland & Karl K. Turekian) 125–156 (Elsevier, 2014).

12. Bais, H. P., Weir, T. L., Perry, L. G., Gilroy, S. & Vicanco, J. M. The role of root exudates in rhizosphere interactions with plants and other organisms. Annu Rev Plant Biol 57, 233–266, doi:10.1146/ (2006).

13. Ettinger, C. L., Williams, S. L., Abbott, J. M., Stachowicz, J. J. & Eisen, J. A. Microbiome succession during ammonification in eelgrass bed sediments. PeerJ 5, e3674 (2017).

14. Trevathan-Tackett, S. M. et al. Sediment anoxia limits microbial-driven seagrass carbon remineralization under warming conditions. FEMS Microbiol Ecol 93 (2017).

15. Drew, E. A. Sugars, cyclitols and seagrass phylogeny. Aquat Bot 15, 387–408 (1983).

16. Pirc, H. Seasonal Changes in Soluble Carbohydrates, Starch, and Energy Content in Mediterranean Seagrasses. Mar Ecol 10, 97–105 (1989).

17. Burdige, D. J., Skoog, A. & Gardner, K. Dissolved and particulate carbohydrates in contrasting marine sediments. Geochim Cosmochim Acta 64, 1029–1041 (2000).

18. Kirchman, D. L. et al. Glucose fluxes and concentrations of dissolved combined neutral sugars (polysaccharides) in the Ross Sea and Polar Front Zone, Antarctica. Deep Sea Res. Part II Top Stud Ocean 48, 4179–4197(2001).

19. Wright, R. R. & Hobbie, J. E. Use of glucose and acetate by bacteria and algae in aquatic ecosystems. Ecology 47, 447–464 (1966).

20. Decker, E. M., Klein, C., Schwindt, D. & von Ohle, C. Metabolic activity of Streptococcus mutans biofilms and gene expression during exposure to xylitol and sucrose. Int J Oral Sci 6, 195–204 (2014).

21. Tringe, S. G. et al. Comparative metagenomics of microbial communities. Sci 308, 554–557 (2005).

22. Tian, L., Dell, E. & Shi, W. Chemical composition of dissolved organic matter in agroecosystems: Correlations with soil enzyme activity and carbon and nitrogen mineralization. Appl Soil Ecol 46, 426–435 (2010).

23. Lievens, B. et al. Microbiology of sugar-rich environments: diversity, ecology and system constraints. Environ Microbiol 17, 278–298 (2015).

24. Moriarty, D. J. W., Iverson, R. L. & Pollard, P. C. Exudation of organic carbon by the seagrass *Halodule wrightii* Aschers. and its effect on bacterial growth in the sediment. J Exp Mar Biol Ecol 96, 115–126 (1986).

25. Holmer, M., Duarte, C. M., Boschker, H. T. S. & Barrón, C. Carbon cycling and bacterial carbon sources in pristine and impacted Mediterranean seagrass sediments. Aquat Microb Ecol 36, 227–237 (2004).

26. Pirc, H. Growth dynamics in *Posidonia oceanica* (L.) Delile. Mar Ecol 6, 141–165 (1985).

27. Danovaro, R. Detritus-bacteria-meiofauna interactions in a seagarss bed *Posidonia oceanica* of the NW Mediterranean. Mar Biol 127, 1–13 (1996).

28. Pirc, H. Seasonal aspects of photosynthesis *in Posidonia oceanica*: Influence of depth, temperature and light intensity. Aquat Bot 26, 203–212 (1986).

29. Koopmans, D., Holtappels, M., Chennu, A., Weber, M. & de Beer, D. High net primary production of Mediterranean seagrass (*Posidonia oceanica*) meadows determined with aquatic eddy covariance. Front Mar Sci 7, 118 (2020).

30. Champenois, W. & Borges, A. V. Seasonal and interannual variations of community metabolism rates of a *Posidonia oceanica* seagrass meadow. Limnol Oceanogr 57, 347–361(2012).

31. Procaccini, G. et al. Depth-specific fluctuations of gene expression and protein abundance modulate the photophysiology in the seagrass *Posidonia oceanica*. Sci Rep 7, 42890 (2017).

32. Hennion, N. et al. Sugars en route to the roots: Transport, metabolism and storage within plant roots and towards microorganisms of the rhizosphere. Physiol Plant 165, 44–57, (2019).

33. Mateo, M. A., Cebrian, J., Dunton, K. H. & Mutchler, T. in Seagrasses: Biology, Ecology and Conservation (eds Anthony W.D. Larkum, R. J. Orth, & C. M. Duarte) 159–192 (Springer, 2006).

34. Holmer, M., Duarte, C. M. & Marbá, N. Sulfur cycling and seagrass (*Posidonia oceanica*) status in carbonate sediments. Biogeochemistry 66, 223–239 (2003).

35. Martin, S. A. & Akin, D. E. Effect of phenolic monomers on the growth and beta-glucosidase activity of *Bacteroides ruminicola* and on the carboxymethylcellulase, beta-glucosidase, and xylanase activities of Bacteroides succinogenes. Appl Environ Microbiol 54, 3019 (1988).

36. Osterholz, H., Kirchman, D. L., Niggemann, J. & Dittmar, T. Environmental drivers of dissolved organic matter molecular composition in the Delaware estuary. Front Earth Sci 4, 95 (2016).

37. Seidel, M. et al. Molecular-level changes of dissolved organic matter along the Amazon river-to-ocean continuum. Mar Chem 177, 218–231 (2015).

38. Zapata, O. & McMillan, C. Phenolic acids in seagrasses. Aquat Bot 7, 307–317 (1979).

39. Agostini, S., Desjobert, J.-M. & Pergent, G. Distribution of phenolic compounds in the seagrass *Posidonia oceanica*. Phytochemistry 48, 611–617 (1998).

40. Hättenschwiler, S. & Vitousek, P. M. The role of polyphenols in terrestrial ecosystem nutrient cycling. Trends Ecol Evol 15, 238–243 (2000).

41. Maie, N., Jaffé, R., Miyoshi, T. & Childers, D. L. Quantitative and qualitative aspects of dissolved organic carbon leached from senescent plants in an oligotrophic wetland. Biogeochemistry 78, 285–314 (2006).

42. Zhalnina, K. et al. Dynamic root exudate chemistry and microbial substrate preferences drive patterns in rhizosphere microbial community assembly. Nat Microbiol 3 (2018).

43. Zwetsloot, M. J., Kessler, A. & Bauerle, T. L. Phenolic root exudate and tissue compounds vary widely among temperate forest tree species and have contrasting effects on soil microbial respiration. New Phytol 218, 530–541 (2018).

44. Zwetsloot, M. J. et al. Prevalent root-derived phenolics drive shifts in microbial community composition and prime decomposition in forest soil. Soil Biol Biochem 145, 107797 (2020).

45. Fitzpatrick, C. R. et al. The Plant Microbiome: From ecology to reductionism and beyond. Annu Rev Microbiol 74, 81–100 (2020).

46. Peiffer, J. A. et al. Diversity and heritability of the maize rhizosphere microbiome under field conditions. Proc Natl Acad Sci U S A 110, 6548 (2013).

47. Bowers, R. M. et al. Minimum information about a single amplified genome (MISAG) and a metagenome-assembled genome (MIMAG) of bacteria and archaea. Nat Biotechnol 35, 725–731 (2017).

48. O’Banion, B. S., O’Neal, L., Alexandre, G. & Lebeis, S. L. Bridging the gap between single-strain and community-level plant-microbe chemical interactions. Mol Plant Microbe Itneract 33, 124–134 (2019).

49. Voges, M. J. E. E. E., Bai, Y., Schulze-Lefert, P. & Sattely, E. S. Plant-derived coumarins shape the composition of an *Arabidopsis* synthetic root microbiome. Proc Natl Acad Sci 116, 12558 (2019).

50. Krell, T. et al. Responses of *Pseudomonas putida* to toxic aromatic carbon sources. J Biotechnol 160, 25–32 (2012).

51. Olsen, J. L. et al. The genome of the seagrass *Zostera marina* reveals angiosperm adaptation to the sea. Nature 530, 331–335 (2016).

52. Calleja, M. L., Marbà, N. & Duarte, C. M. The relationship between seagrass (*Posidonia oceanica*) decline and sulfide porewater concentration in carbonate sediments. Estuar Coast Shelf Sci 73, 583–588 (2007).

53. Ramm, A. E. & Bella, D. A. Sulfide production in anaerobic microcosms. Limnol Oceanogr 19, 110–118 (1974).

54. Holmer, M., Andersen, F. Ø., Nielsen, S. L. & Boschker, H. T. S. The importance of mineralization based on sulfate reduction for nutrient regeneration in tropical seagrass sediments. Aquat Bot 71, 1–17 (2001).

55. Waycott, M. et al. Accelerating loss of seagrasses across the globe threatens coastal ecosystems. Proc Natl Acad Sci U S A 106, 12377–12381(2009).

56. Sogin, E. M., Puskas, E., Dubilier, N. & Liebeke, M. Marine metabolomics: a method for the non-targeted measurement of metabolites in seawater by gas-chromatography mass spectrometry. mSystems, 4, e00638–19 (2020).

57. Lisec, J., Schauer, N., Kopka, J., Willmitzer, L. & Fernie, A. R. Gas chromatography mass spectrometry-based metabolite profiling in plants. Nat Protoc 1, 387–396 (2006).

58. Liebeke, M. & Puskas, E. Drying enhances signal intensities for global GC-MS metabolomics. Metabolites 9, (2019).

59. Holman, J. D., Tabb, D. L. & Mallick, P. Employing ProteoWizard to convert raw mass spectrometry data. Curr Protoc Bioinformatics 46, 11–19 (2014).

60. Smith, C. A., Want, E. J., O’Maille, G., Abagyan, R. & Siuzdak, G. XCMS: Processing mass spectrometry data for metabolite profiling using nonlinear peak alignment, matching, and identification. Anal Chem, 78, 779–787 (2006).

61. Kuhl, C., Tautenhahn, R., Bottcher, C., Larson, T. R. & Neumann, S. CAMERA: an integrated strategy for compound spectra extraction and annotation of liquid chromatography/mass spectrometry data sets. Analytical Chemistry 84, 283–289 (2012).

62. Dittmar, T., Koch, B., Hertkorn, N. & Kattner, G. A simple and efficient method for the solid-phase extraction of dissolved organic matter (SPE-DOM) from seawater. Limnol Oceanogr 6, 230–235 (2008).

63. Seidel, M. et al. Biogeochemistry of dissolved organic matter in an anoxic intertidal creek bank. Geochim Cosmochim Acta 140, 418–434 (2014).

64. Riedel, T. & Dittmar, T. A method detection limit for the analysis of natural organic matter via Fourier transform ion cyclotron resonance mass spectrometry. Anal Chem 86, 8376–8382 (2014).

65. Koch, B. P. & Dittmar, T. From mass to structure: an aromaticity index for high-resolution mass data of natural organic matter. Rapid Commun in Mass Spectrom 20, 926–932 (2006).

66. Weber, M. et al. In situ applications of a new diver-operated motorized microsensor profiler. Environ Sci Technol 41, 6210–6215 (2007).

67. Martin, B. C. et al. Oxygen loss from seagrass roots coincides with colonisation of sulphide-oxidising cable bacteria and reduces sulphide stress. ISME J 13, 707–719 (2019).

68. Micheline, G.-D. & Bernadette, R. Phenolic fingerprint of the seagrass *Posidonia oceanica* from four locations in the Mediterranean Sea: first evidence for the large predominance of chicoric acid. Bot Mar 58, 379–391(2015).

69. Grignon-Dubois, M. & Rezzonico, B. in Botanica Marina Vol. 58 379 (2015).

70. Marta E. Torres, Mix, A. C. & Rugh, W. D. Precise δ^13^C analysis of dissolved inorganic carbon in natural waters using automated headspace sampling and continuous-flow mass spectrometry. Limnol Oceanogr 3, 349–360 (2005).

71. Torti, A., Jorgensen, B. B. & Lever, M. A. Preservation of microbial DNA in marine sediments: insights from extracellular DNA pools. Environ Microbiol 20, 4526–4542 (2018).

72. Zhou, J., Bruns, M. A. & Tiedje, J. M. DNA Recovery from soils of diverse composition. Appl Environ Microbiol 62, 316–322 (1996).

73. Lever, M. A. et al. A modular method for the extraction of DNA and RNA, and the separation of DNA pools from diverse environmental sample types. Front Microbiol 6, 476 (2015).

74. Ansede, J. H., Friedman, R. & Yoch, D. C. Phylogenetic analysis of culturable dimethyl sulfide-producing bacteria from *Spartina* dominated salt marsh and estuarine water. Appl Environ Microbiol 67, 1210 (2001).

75. Callahan, B. J. et al. DADA2: High-resolution sample inference from Illumina amplicon data. Nat Methods 13, 581–583 (2016).

76. McMurdie, P. J. & Holmes, S. phyloseq: an R package for reproducible interactive analysis and graphics of microbiome census data. PLoS One 8, e61217 (2013).

77. Gruber-Vodicka, H. R., Seah, B. K. B. & Pruesse, E. phyloFlash – Rapid small subunit rRNA profiling and targeted assembly from metagenomes. mSystems, 5:e00920–20 (2020).

78. Quast, C. et al. The SILVA ribosomal RNA gene database project: improved data processing and web-based tools. Nucleic Acids Res 41, D590–D596 (2013).

79. Nurk, S., Meleshko, D., Korobeynikov, A. & Pevzner, P. A. metaSPAdes: a new versatile metagenomic assembler. Genome Res 27, 824–834 (2017).

80. Li, D., Liu, C. M., Luo, R., Sadakane, K. & Lam, T. W. MEGAHIT: an ultra-fast single-node solution for large and complex metagenomics assembly via succinct de Bruijn graph. Bioinformatics 31, 1674–1676 (2015).

81. Kang, D. D., Froula, J., Egan, R. & Wang, Z. MetaBAT, an efficient tool for accurately reconstructing single genomes from complex microbial communities. PeerJ 3, e1165 (2015).

82. Alneberg, J. et al. Binning metagenomic contigs by coverage and composition. Nat Methods 11, 1144–1146 (2014).

83. Wu, Y.-W., Simmons, B. A. & Singer, S. W. MaxBin 2.0: an automated binning algorithm to recover genomes from multiple metagenomic datasets. Bioinformatics 32, 605–607 (2015).

84. Sieber, C. M. K. et al. Recovery of genomes from metagenomes via a dereplication, aggregation and scoring strategy. Nat Microbiol 3, 836–843 (2018).

85. Parks, D. H., Imelfort, M., Skennerton, C. T., Hugenholtz, P. & Tyson, G. W. CheckM: assessing the quality of microbial genomes recovered from isolates, single cells, and metagenomes. Genome Res 25, 1043–1055 (2015).

86. Parks, D. H. et al. A standardized bacterial taxonomy based on genome phylogeny substantially revises the tree of life. Nat Biotechnol 36, 996–1004 (2018).

87. Hyatt, D. et al. Prodigal: prokaryotic gene recognition and translation initiation site identification. BMC Bioinform 11, 119 (2010).

88. Zhang, H. et al. dbCAN2: a meta server for automated carbohydrate-active enzyme annotation. Nucleic Acids Res 46, W95–W101 (2018).

89. Buchfink, B., Xie, C. & Huson, D. H. Fast and sensitive protein alignment using DIAMOND. Nat Methods 12, 59–60 (2015).

90. Magrane, M. & Consortium, U. UniProt Knowledgebase: a hub of integrated protein data. Database 2011 (2011).

91. Jones, P. et al. InterProScan 5: genome-scale protein function classification. Bioinformatics 30, 1236–1240 (2014).

92. Finn, R. D. et al. Pfam: the protein families database. Nucleic Acids Res 42, D222–D230 (2013).

93. Käll, L., Krogh, A. & Sonnhammer, E. L. L. A combined transmembrane topology and signal peptide prediction method. Journal of Molecular Biology 338, 1027–1036 (2004).

94. Huerta-Cepas, J. et al. eggNOG 5.0: a hierarchical, functionally and phylogenetically annotated orthology resource based on 5090 organisms and 2502 viruses. Nucleic Acids Res 47, D309–D314 (2019).

95. Kopylova, E., Noé, L. & Touzet, H. SortMeRNA: fast and accurate filtering of ribosomal RNAs in metatranscriptomic data. Bioinformatics 28, 3211–3217 (2012).

96. Bray, N. L., Pimentel, H., Melsted, P. & Pachter, L. Near-optimal probabilistic RNA-seq quantification. Nat Biotechnol 34, 525–527 (2016).

97. Lee, M. D. GToTree: a user-friendly workflow for phylogenomics. Bioinformatics, 15, 4162–4164 (2019).

98. Eddy, S. Accelerated profile HMM searches. PloS Comput Biol 7, e1002195 (2011).

99. Edgar, R. MUSCLE: a multiple sequence alignment method with reduced time and space complexity. BMC Bioinform 5, 113 (2004).

100. Capella-Gutiérrez, S., Silla-Martínez, J. & Gabaldón, T. TrimAl: a tool for automatic alignment trimming. Bioinformatics 25, 1972–1973 (2009).

101. Nguyen, L., Schmidt, H., Haeseler, A. v. & Minh, B. IQ-TREE: a fast and effective stochastic algorithm for estimating maximum likelihood phylogenies. Mol Biol Evol 32, 268–274 (2015).

102. Letunic, I. & Bork, P. Interactive Tree Of Life (iTOL) v4: recent updates and new developments Nucleic Acids Res 47, W256–W259 (2019).

103. Diepenbroek, M. et al. in Informatik 2014 (eds E. Plödereder, L. Grunske, E. Schneider, & D. Ull) 1711–1721 (Gesellschaft für Informatik e.V., 2014).

104. Yilmaz, P. et al. Minimum information about a marker gene sequence (MIMARKS) and minimum information about any (x) sequence (MIxS) specifications. Nat Biotech 29, 415–420 (2011).

