## Supporting Information for "Sugars dominate the seagrass rhizosphere"

1- Max Planck Institute for Marine Microbiology
2- University of California at Merced
3- University of Vienna
4- University of Oldenburg
5- Stazione Zoologica Anton Dohrn

### Supporting Information

Supporting tables:

Table S1. Average concentrations of sugars across sampling sites, seagrass species, seasons, times and depths

Table S2. Sugar and dissolved organic carbon concentrations, results from statistical tests

Table S3. Incubation rates of sucrose degradation from seagrass sediments

Table S4. Metagenomic library statistics and accession numbers

Table S5. Metagenomic bin statistics

Table S6. Metatranscriptomic expression data for sucrose specialist MAGs

Table S7. Description of mass spectrometry imaging datasets

***Zostera marina.*** We observed less then 2 µM of sucrose (**Table S1**) underneath *Z. marina* meadows in the Baltic sea, which is three orders of magnitude lower than the maximum sucrose concentrations observed underneath the Mediterranean and Caribbean seagrass meadows investigated in this study (**Table S1**). There are likely many environmental and biological factors that can explain our observations, of which we highlight three below. First, low sugar concentrations in *Z. marina* sediments is likely related to the lower availability of light in the Baltic Sea in comparison to the Mediterranean or Caribbean sampling sites. Under periods of low light, *Z. marina* produces less photosynthate, resulting in a decrease of both non-structural carbohydrates, including sucrose, and polyphenolic compounds^1,2^. Consequently, we predict under the low light regime in the Baltic Sea, *Z. marina* does not have sufficient reserves of sucrose or polyphenolic compounds to excrete to its rhizospheres. Second, salinity is also correlated to sucrose content in seagrass tissues, where seagrasses living in lower salinity waters have reduced sucrose concentrations in their tissues^3^. The low salinity of the Baltic Sea sampling site in Kiel Bay of ca. 20‰^4^ could alter the osmotic balance in *Z. marina*, which would reduce the amount of sugars in its tissues, and consequently the amount of sugars it excretes. Finally, we predict nutrients also play a major role in sucrose concentrations underneath *Z. marina* meadows. Specifically, higher nutrients in the Baltic Sea lead to increased growth of epiphytic algae on *Z. marina* leaves, which decrease seagrass productivity^5^ and consequently sediment sucrose concentrations.

**Sediment incubations**. To examine if seagrass-derived phenolics inhibit microbial respiration of sucrose, we incubated sediments from a Mediterranean *P. oceanica* meadow with labeled ^13^C_12_-sucrose in the presence and absence of oxygen as well as seagrass-derived phenolics. The volumetric rates determined during these incubations were converted to areal rates assuming an active layer of 20 cm to compare them to total ecosystem productivity and respiration rates reported elsewhere^6^. Under oxic conditions, sediments collected in the upper 3 cm depth showed similar removal rates of sucrose across all treatments at 86 mmol(C)/m²d^-1^. The rates of ^13^CO_2_ released corresponded well with removal rates, and ranged between 64 and 83 mmol(C)/m²d^-1^ (**Table S3**). These experimental rates are unlikely to be significant in situ since the concentration of sucrose close to the sediment surface is low at an average of 0.35 µM. Under oxic conditions, we assume the phenolics were quickly oxidized and removed from the incubation, thereby allowing the microbial community from inside the meadow to respire sucrose (**Figure 3**).

Sediment collected from the anoxic layers underneath the meadow and diluted 15-fold with anoxic seawater, removed sucrose at a rate corresponding to 1891 mmol/m²d^-1^, and respired sucrose fully to CO_2_ at a rate of 232 mmol(C)/m²d^-1^. These rates would not be supported in the environment by a *P. oceanica* meadow, as they are higher than the gross primary production (GPP) of the meadow in the highly productive month May (average 169.2 mmol(C)/m²d^-1^)^6^. Flushing the sediment with five volumes of artificial, anoxic seawater before the incubation (to remove seagrass-derived phenolics), reduced the rates slightly to 1589 mmol(C)/m²d^-1^ and 158 mmol(C)/m²d^-1^, which is still much higher than what could be supported in the environment. Since we consistently detected high concentrations of sucrose under the meadows, there must be an inhibitory effect in the environment.

**Ecosystem metabolism.** Potential rates of sucrose respiration from our incubations were compared to previously published oxygen fluxes measured on a *P. oceanica* seagrass meadow in Sant’Andra Bay, Elba, Italy with the eddy correlation technique^6^. First, net ecosystem metabolism was derived from oxygen fluxes assuming that oxygen is a proxy for mineralization and primary production following a ratio 1 mol CO_2_:1 mol O_2_ (for a review of oxygen dynamics in marine sediments see^7^). For *P. oceanica* this yields a gross primary production of 162 mmol(C) m^-2^d^-1^ (GPP), a respiration of 85 mmol(C) m^-2^d^-1^ (R), and a net ecosystem metabolism of 77 mmol(C) m^-2^d^-1^ (NEM). The sucrose degradation rates in absence of phenolics ranged between 64 and 232 mmol(C) m^-2^d^-1^, therefore they exceed the actual overall ecosystem respiration R and cannot be explained by the NEM. In contrast, the sucrose respiration in the presence of phenolics reduces to 20 mmol(C) m^-2^d^-1^ and is likely included in the respiration measurement R. Therefore, under the assumption that our measured sucrose respiration is mostly driven by excess primary production of the plant, the sucrose degradation can be directly compared to the NEM: 20 mmol(C) m^-2^d^-1^ / 77 mmol(C) m^-2^d^-1^ = 29 %. However, as we cannot exclude that the plant is excreting more sucrose then the amount that is respired, the 29 % represents rather the lower limit.

**General metabolism of sucrose specialists.** To determine the metabolism of the six sucrose specialists, we looked for the expression of key marker genes involved in select metabolic pathways. Because our MAGs were incomplete (72.8% to 94.8%), it is possible some metabolic genes are missing from the MAGs.

Both MAGs 76 and 154 lacked a cytochrome c terminal oxidase, making them likely anaerobes as the MAGs do not have the genes needed for oxygen respiration. MAG 154 (Desulfosarcinaceae sp.) contained genes for sulfate reduction, hydrogen oxidation (hydrogenase large and small subunit; EC 1.12.99.6), and nitrogen fixation (nitrogenase proteins, NifA, NifB, NifE, and NifH), all of which were expressed. MAG 154 was more abundant inside the meadow then at the edge or outside (**Figure S8**). We predict this population of bacteria is capable of fermenting plant-based sugars based on the expression patterns of multiple types of glycoside hydrolase enzymes. MAG 76 (Beggiatoales sp.) contained genes for sulfur oxidation (SoxB,SoxX, SoxY, and SoxZ) and genes that use nitrate an electron acceptor (respiratory nitrate reductase, EC 1.7.99.4). It also had genes for hydrogen oxidation (hydrogenase large and small subunit; EC 1.12.99.6) and nitrogen fixation (nitrogenase beta and alpha chains; EC 1.18.6.1). These genes were also expressed in our sediment metatranscriptomes.

The remaining four MAGs contained genes for oxygen respiration which were expressed in the transcriptomic libraries. Expression data indicated MAG 207 (Thiohalomonadales sp.) fixed CO_2_ through the Calvin Cycle (RuBisCO, EC 4.1.1.39), oxidized reduced sulfur compounds (SoxA, SoxB, SoxY, SoxZ) and possibly hydrogen (hydrogenase, EC 1.12.99.6), reduced nitrite to nitrous oxide (Nitric-oxide reductase, EC 1.7.99.7) and fixes nitrogen (nitrogenase; NifA, NifB; nitric-oxid reductase EC 1.7.99.7).

MAG 142 (Xanthomonadales) also contained genes for autotrophic fixation of CO_2_ through the Calvin cycle (EC 4.1.1.39) and hydrogen oxidation (hydrogenase, EC 1.12.99.6), both of which were expressed in the sediment metatranscriptomes. MAG 209 (Verrucomicrobiales) and 438 (Gammaproteobacteria sp.) contained genes for the general uptake of organic material through heterotrophy.

Five out of the six MAGs also contained genes involved in the breakdown of phenolics, many of which were expressed the sediment metatranscriptomes. This is intriguing since many phenolic compounds limit bacterial growth and produce significant antibacterial activity, often accompanied by an increase of reactive oxygen species^8^. Some bacteria respond to the presence of phenolics with the production of antioxidants, like glutathione or enzymes like catalase and dismutase to counterbalance reactive oxygen stress. The sucrose specialists in our study have at least two pathways for defense against phenolics: They expressed genes for phenolic degradation pathways (benzoyl-CoA and beta-ketoadipate pathways) and the production of antioxidant compounds and enzymes (e.g. catalase, superoxide dismutase, and glutathione peroxidase ). Our data suggest the expression of these enzymes enables these taxa to use sucrose, and other sugars, in their metabolism.

**References**

1	Wong, Y. H. *et al.* High-throughput transcriptome sequencing of the cold seep mussel Bathymodiolus platifrons. *Sci Rep* **5**, 16597, doi:10.1038/srep16597 (2015).

2	Teresa, A., Richard, C. Z., Donald, G. K. & Randall, S. A. Resource allocation and sucrose mobilization in light-limited eelgrass Zostera marina. *MEPS* **187**, 121-131 (1999).

3	Salo, T. & Pedersen, M. F. Synergistic effects of altered salinity and temperature on estuarine eelgrass (Zostera marina) seedlings and clonal shoots. *J of Exp Mar Biol Ecol* **457**, 143-150 (2014).

4	Feistel, R. *et al.* Density and Absolute Salinity of the Baltic Sea 2006–2009. *Ocean Sci.* **6**, 3-24, doi:10.5194/os-6-3-2010 (2010).

5	Baden, S., Boström, C., Tobiasson, S., Arponen, H. & Moksnes, P.-O. Relative importance of trophic interactions and nutrient enrichment in seagrass ecosystems: A broad-scale field experiment in the Baltic−Skagerrak area. *Limnol. Ocean* **55**, 1435-1448 (2010).

6	Koopmans, D., Holtappels, M., Chennu, A., Weber, M. & de Beer, D. High Net Primary Production of Mediterranean Seagrass (*Posidonia oceanica*) Meadows Determined With Aquatic Eddy Covariance. *Front Mar Sci* **7**, 118 (2020).

7	Glud, R. N. Oxygen dynamics of marine sediments. *Mar Biol Res* **4**, 243-289 (2008).

8	Kohanski, M. A., Dwyer, D. J., Hayete, B., Lawrence, C. A. & Collins, J. J. A Common Mechanism of Cellular Death Induced by Bactericidal Antibiotics. *Cell* **130**, 797-810 (2007).

**Supporting Figures**

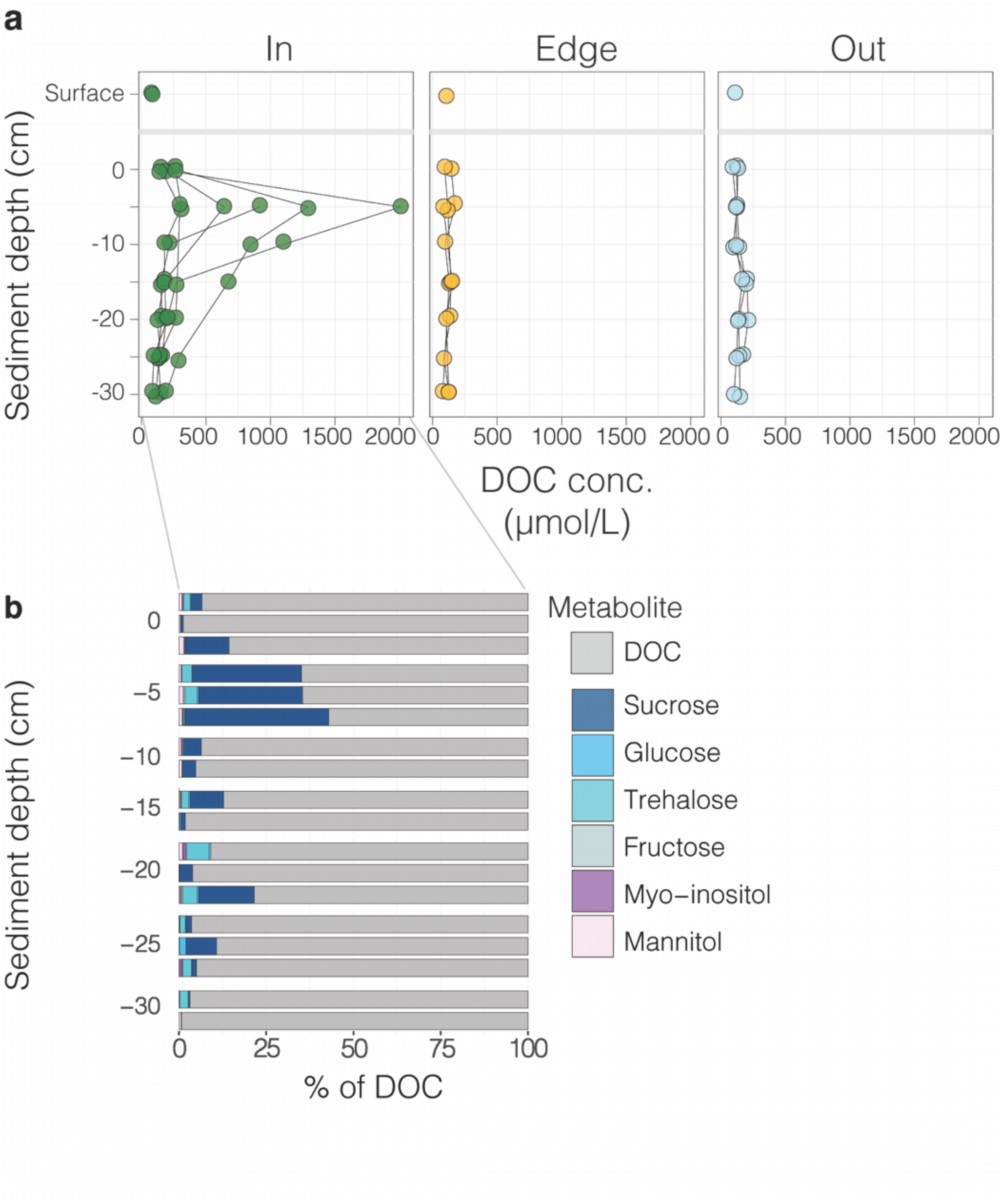

**Figure S1 | Dissolved organic carbon (DOC) concentrations were higher inside the meadow then at the edge or outside.** **a,** DOC concentrations in sediment porewater profiles from inside, at the edge and outside of a *P. oceanica* meadow show DOC concentrations were significantly higher in porewaters inside the meadow (*p* < 0.001; **Table S2**). **b,** A subset of the DOC samples collected inside the meadow show that sugars made up to 40% of the DOC composition within the 5 cm depth, where seagrass roots dominated the sediment.

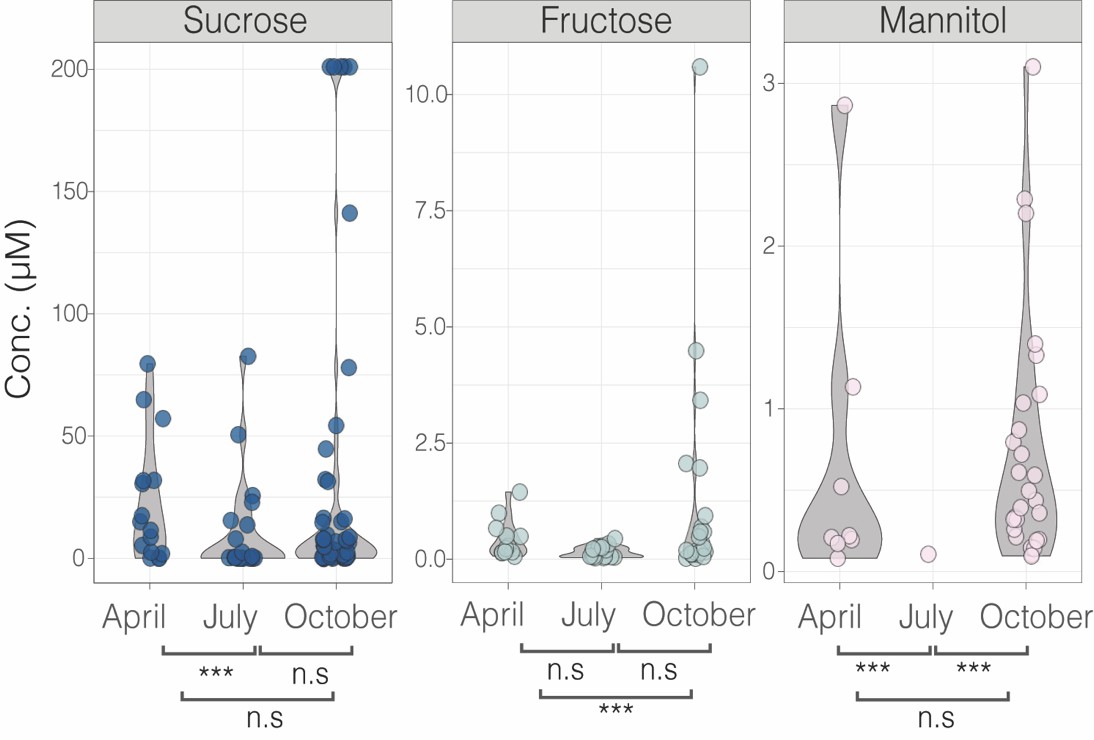

**Figure S2 | Sugar concentrations in the porewater underneath a *P. oceanica* meadow change across seasons.** Violin plots show sugar concentrations varied with sampling seasons. Sugar concentrations were higher in April and October than in July. *** = two-way ANOVA *p-value* < 0.05, n.s.= not significant **(Table S2).**

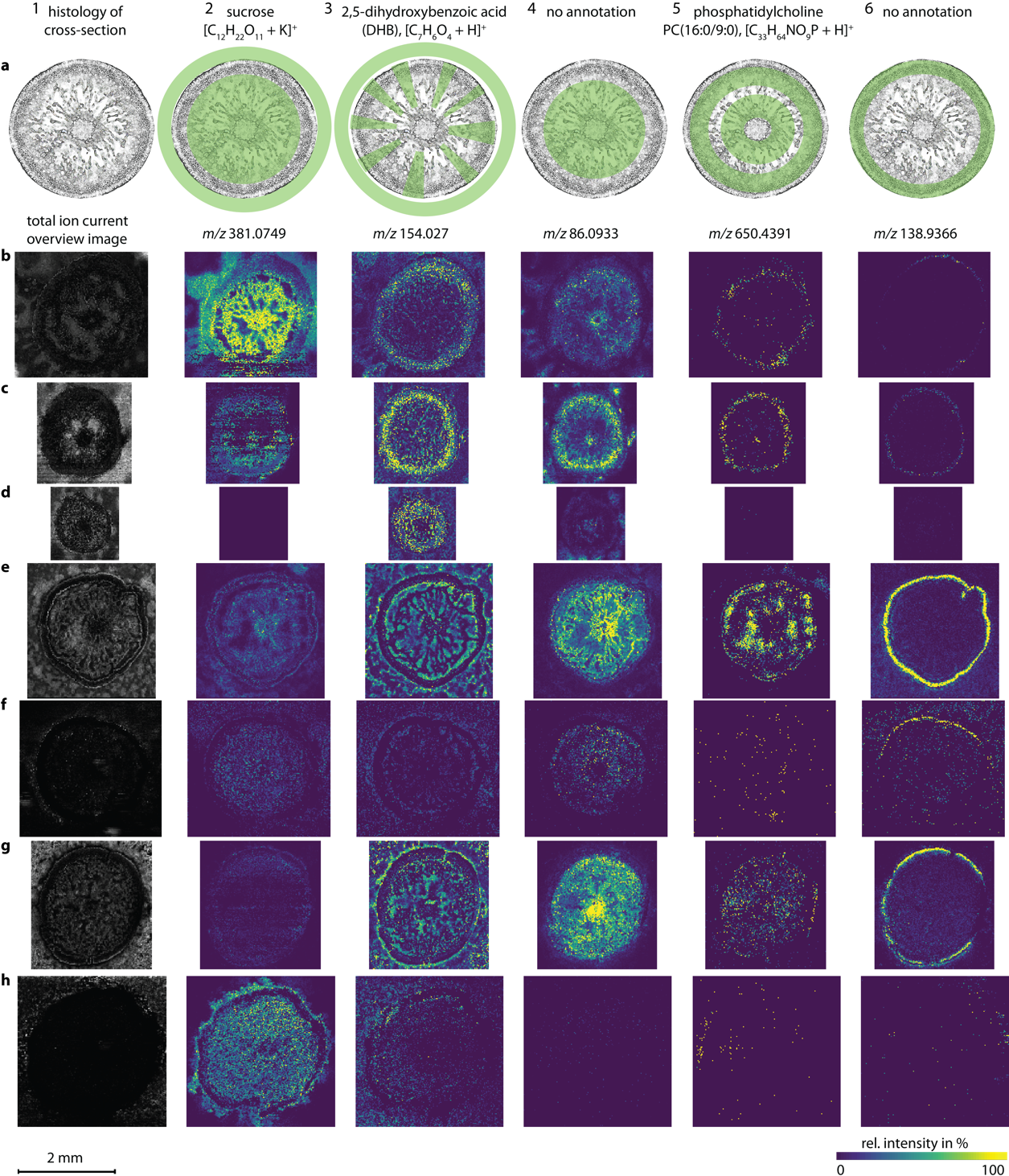

**Figure S3 |** **Distribution of sucrose and other small molecules across seagrass roots.** Replicates and controls for matrix-assisted laser desorption/ionization mass spectrometry imaging in rows **b** to **h**, shown for five different compounds in columns 2 to 6. **a,** Column 1 shows a schematic of a cross section through a seagrass root. Column 2 to 6, row **a** show overlays of the seagrass root and the distribution patterns of different metabolites in green. Samples in rows **b**-**f,** were embedded and cryo-sectioned in carboxymethyl cellulose embedding medium. Samples in rows **g** and **h** were not embedded for cryo-sectioning to test if the sucrose leaked into the aqueous embedding medium. The sucrose at the outer boarder of the root appeared to leak into the embedding medium. Ion images in column 1 show the total ion current (TIC) image for each dataset. Metabolites in column 3 to 6 show different distributions that were consistent across the 7 control datasets. For further descriptions of each dataset see **Table S7**.

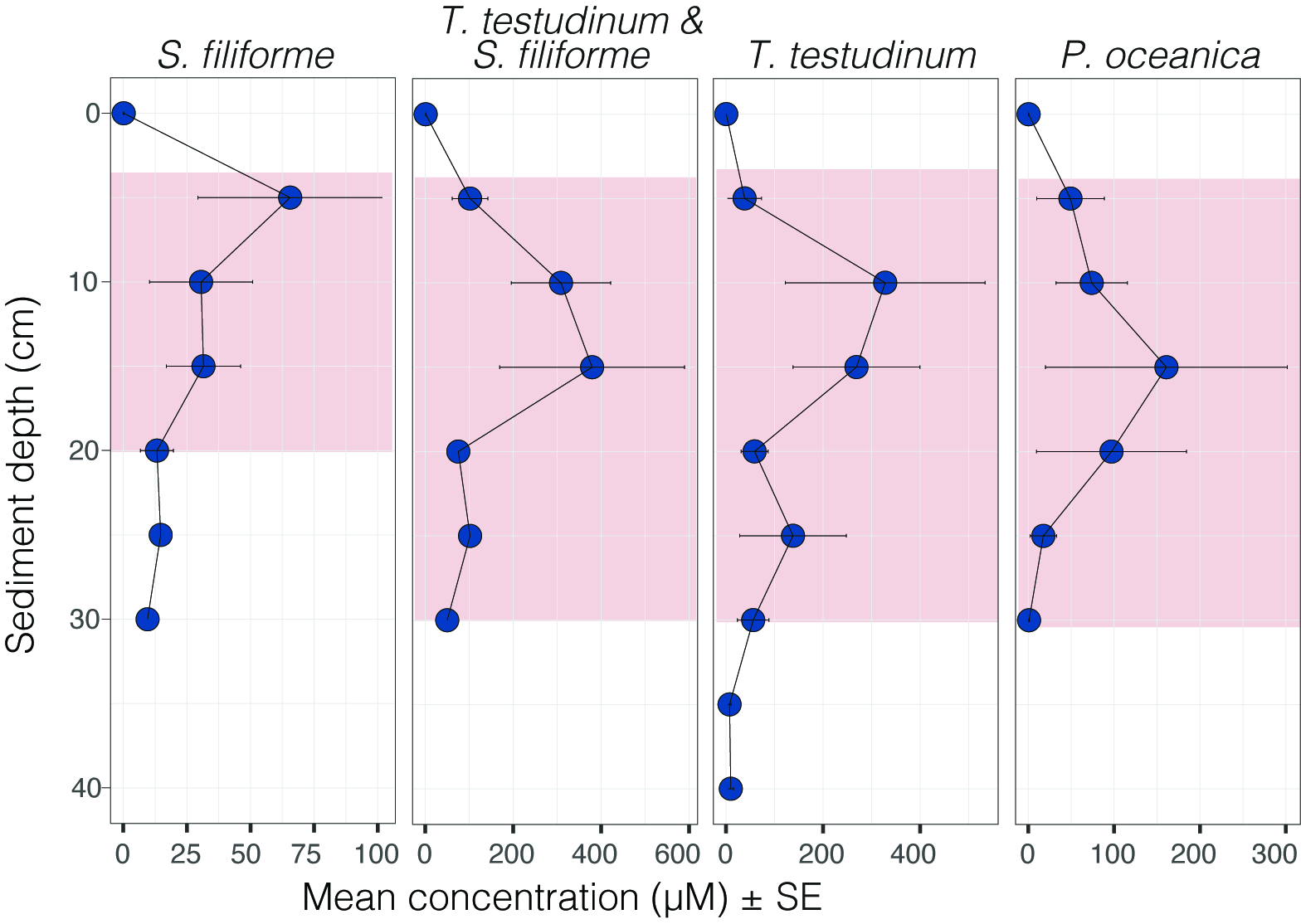

**Figure S4 | Porewater sucrose concentrations are highest at the sediment root interface.** Points represent mean ± standard error of sucrose concentrations (µM)(**Table S1**). Pink boxes show known depth of seagrass roots for each meadow type based on literature reports.

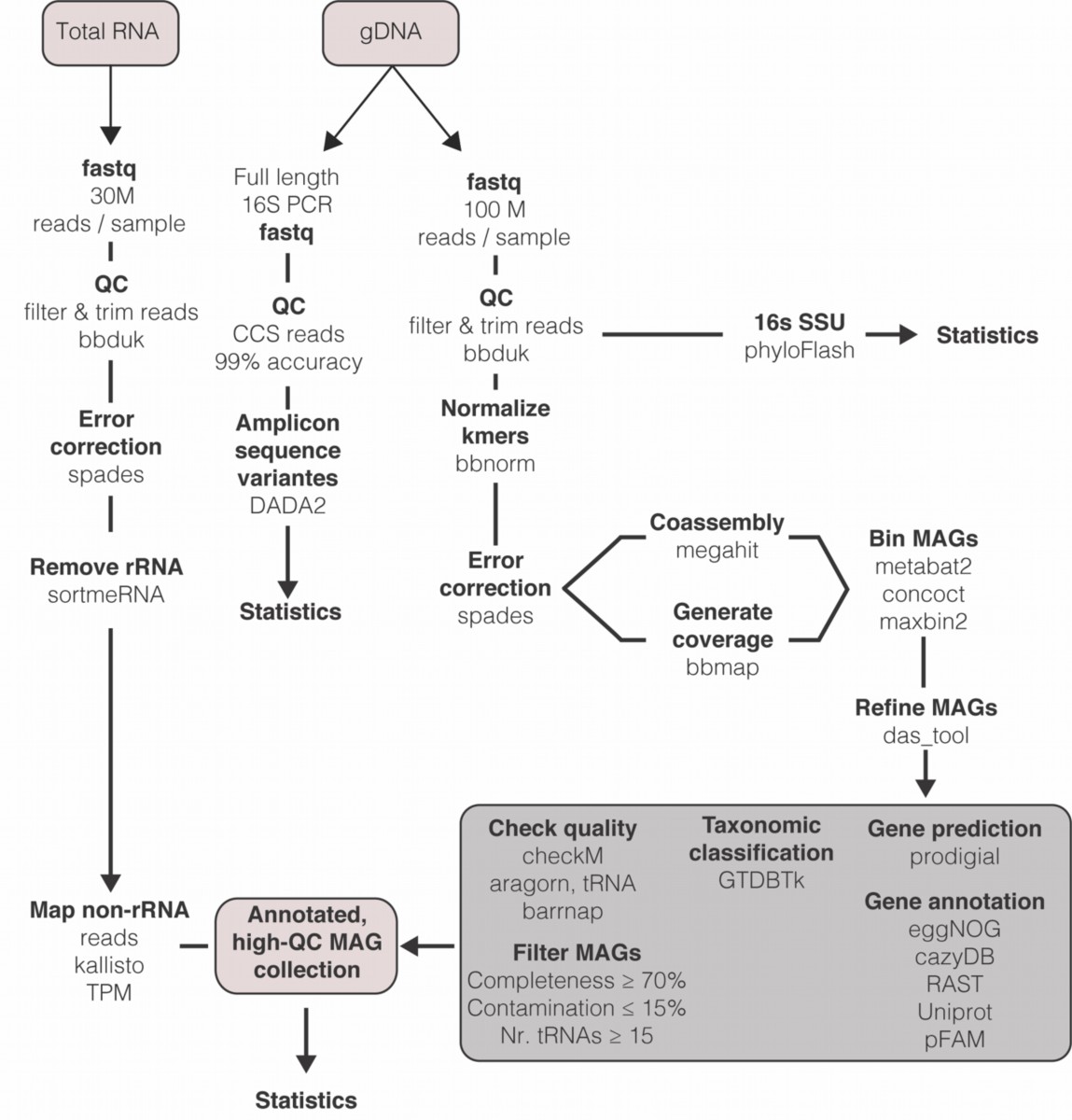

**Figure S5 | Bioinformatic pipeline used to process sediment metagenomes and metatranscriptomes.** Following genomic DNA and total RNA extraction and sequencing, both full length 16S rRNA PacBio data and Illumina metagenomic libraries were used to determine the composition of the microbial community and its metabolism.

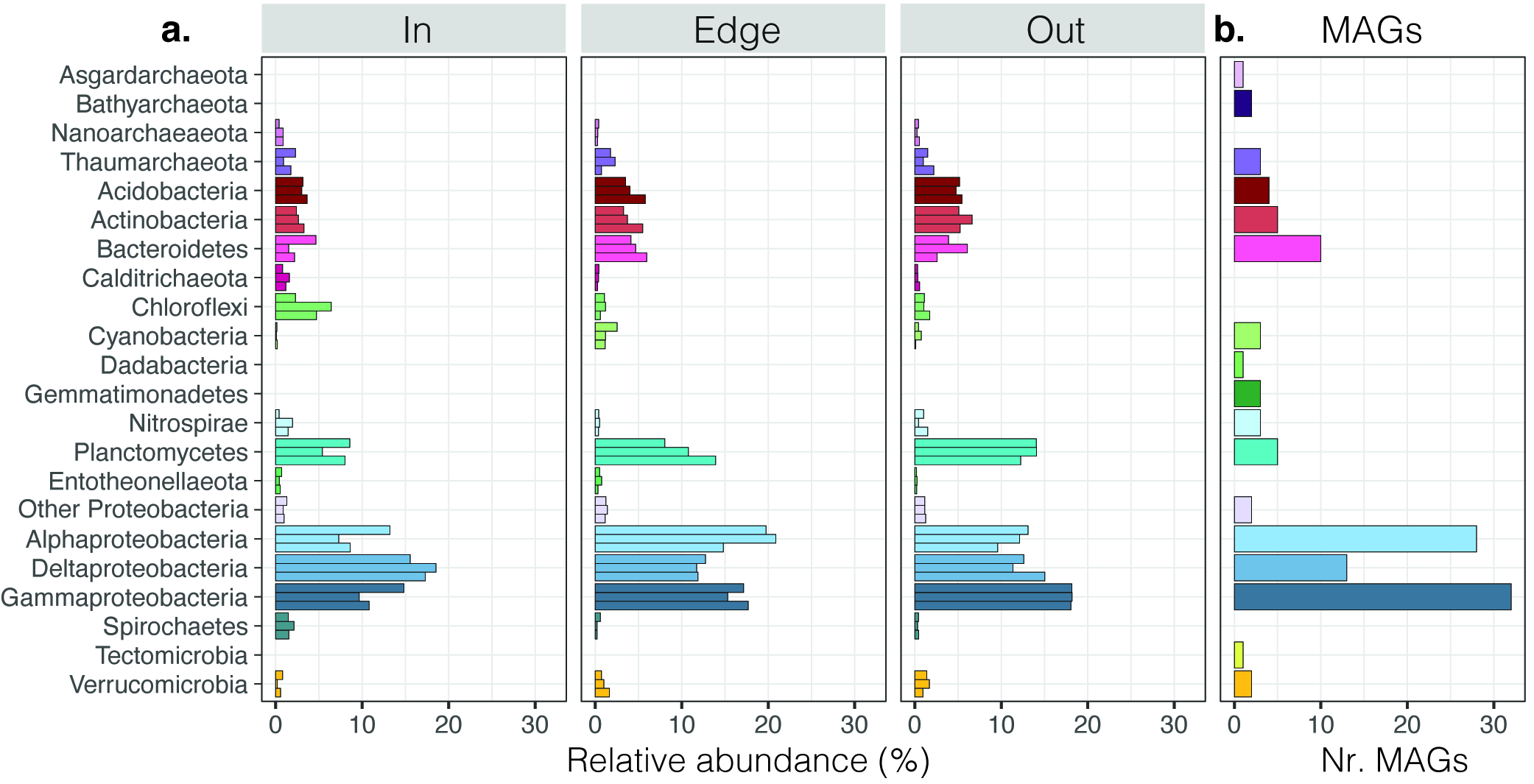

**Figure S6 |** **Composition of sediment microbial communities according to phyla.** **a.** The relative abundance of taxonomic phyla from each metagenomic library collected inside, at the edge and outside a Mediterranean *P. oceanica* meadow off of the island of Elba, Italy. **b.** The number of reconstructed MAGs classified by phylum, obtained from the binned co-assembly of all metagenomic reads from these sediments.

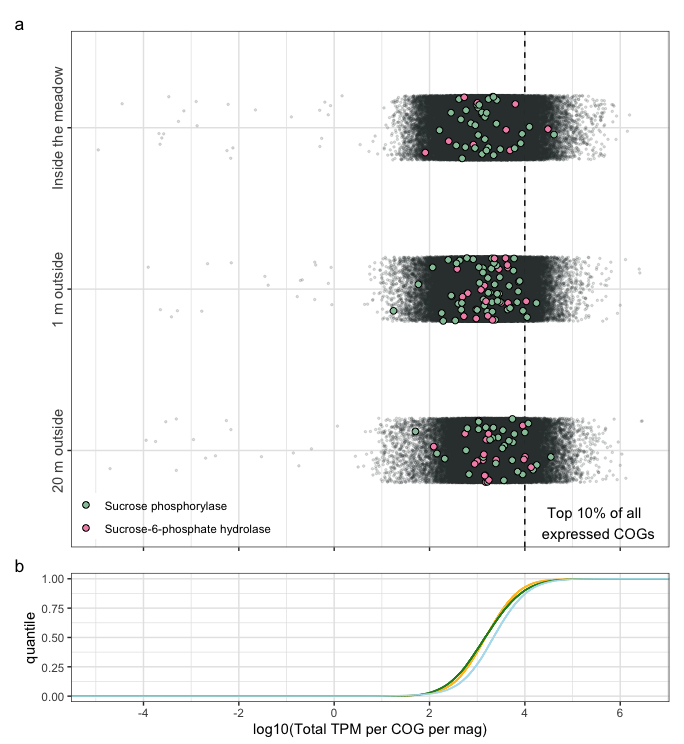

**Figure S7 | Relative expression of sucrose degrading genes is low. a,** The total accumulative expression of clusters of orthologous groups of proteins per MAG from sediments collected inside, at the edge and outside a Mediterranean *P. oceanica* meadow off the island of Elba, Italy. Each point represents the total TPM of a COG category within an individual MAG summed across habitat-specific libraries. COGs predicted to degrade sucrose through either phosphorylase (green) or hydrolase activity (pink) are colored and enlarged to aid visualization. The genes within the top 10% of all expressed COGs fall to the right of the dashed black line as calculated from the cumulative distribution curves (**b**).

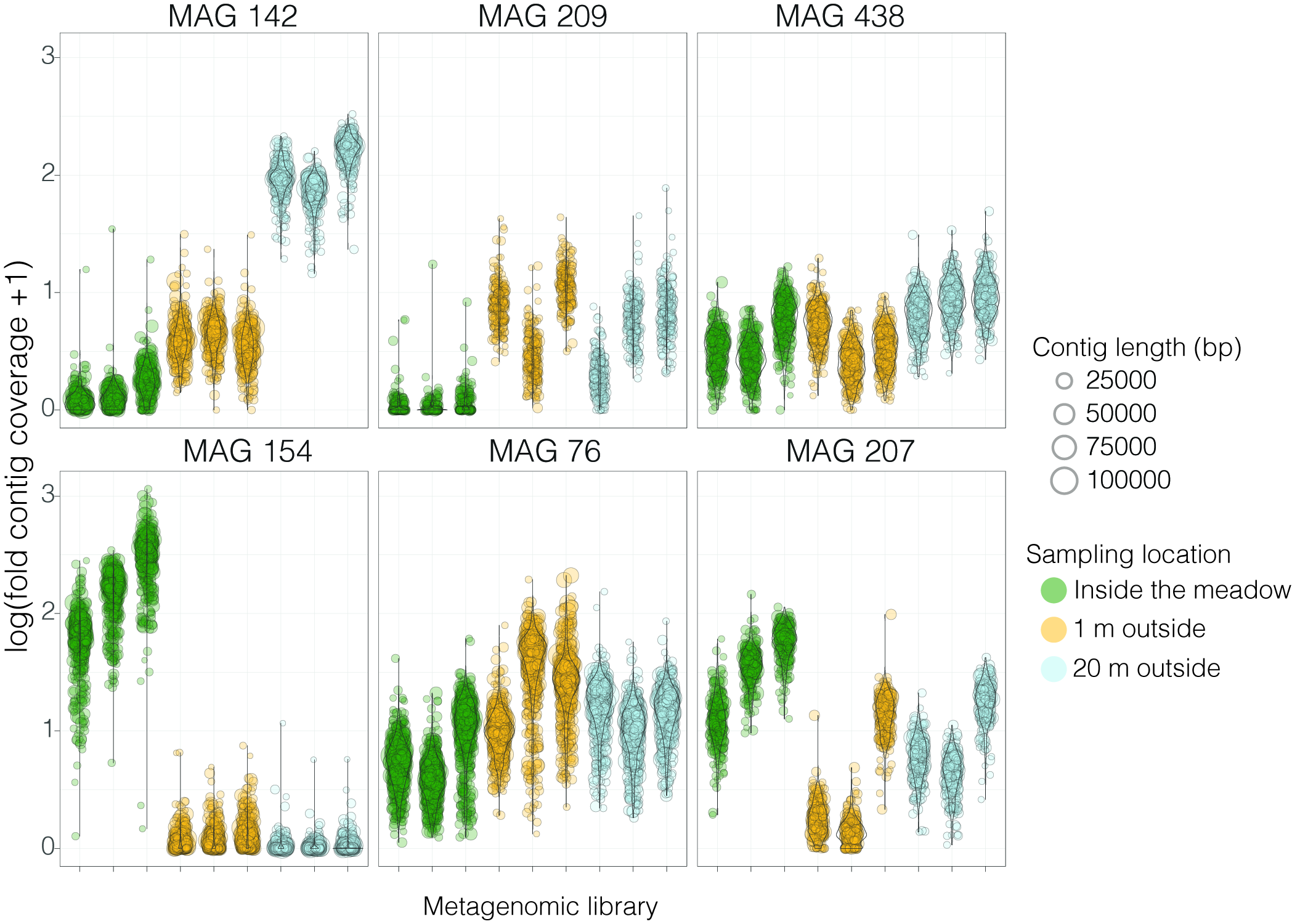

**Figure S8 | Abundance of sucrose specialists across sampling sites.** Abundance plots of sucrose specialists across habitats (In, Edge, Out) based on metagenomic read coverage of individual contigs. Each point represents a contig sized by length (bp).
